# Microtubule architecture and detyrosination bidirectionally modulate sarcomere shortening in skeletal muscle fibers

**DOI:** 10.64898/2026.07.20.739552

**Authors:** Osman Esen, Eva Larose, Leander A. Vonk, Nienke ten Cate, Tyler J. Kirby

## Abstract

Although microtubules (MT) are established regulators of striated muscle mechanics, how MT lattice organization and post-translational modifications (PTM) individually shape contractility in healthy skeletal muscle fibers remains incompletely understood. We used an ex vivo single muscle fiber culture under unloading, examining acetylation and detyrosination (deTyr). During long-term 2D culture, the MT lattice was disrupted by transverse MT depletion without changes in MT abundance. In individual fibers, MT structure, but not abundance, positively correlated with sarcomere shortening non-linearly. When fibers were cultured in 3D hydrogels, the MT lattice was similarly disrupted yet shortening was preserved, with increased longitudinal MTs and decreased deTyr-MTs. Pharmacologically, contractility increased with either a decrease (parthenolide) or an increase (Taxol) in deTyr-MTs, and Taxol further rescued the MT lattice. Our findings identify MT organization and detyrosination, rather than MT abundance, as key determinants of muscle fiber contractility, positioning deTyr-MT as a load-responsive, bidirectional marker relevant to disuse atrophy and aging.

## Introduction

Skeletal muscle constitutes approximately 40% of body mass and is crucial for several tasks including force production, thermogenesis, and protection of vital organs (1,2). Its function depends on a hierarchical structure composed of muscle fibers, myofibrils, and sarcomeres, the smallest units that can contract (1–5). To preserve structural integrity and force transmission, muscle fibers rely on a network of non-sarcomeric cytoskeletal components, including actin filaments, microtubules (MTs), and intermediate filaments (2,3,6,7). Together, this non-sarcomeric cytoskeletal network maintains cellular integrity (6,8), provides mechanical strength (9,10), mediates intracellular force transmission (mechanotransduction) (11), anchors cellular organelles (i.e. mitochondria and nuclei) (12), and links the entire contractile apparatus to the broader mechanical framework of muscle fibers (3,13,14). Each of the non-sarcomeric cytoskeletal elements plays a unique yet cooperative role within the structural framework of muscle fibers (8,12,15), whereby disruption of this intricate network can impair contraction and is associated with various myopathies (3,5,6,13,16). A few examples include desminopathies and dystrophinopathies, which feature sarcomere misalignments (5,17–19) and disorganized MT networks (13,17,19,20), highlighting the importance of an intact cytoskeleton for effective muscle function (21).

During myogenesis, MTs act as a framework to guide myoblasts in forming myotubes and eventually mature contracting muscle fibers (7,19,22). The MTs reorganize from centrosome-nucleated arrays in myoblasts into a mesh-like network, known as MT lattices, composed of intersecting longitudinal and transverse filament populations with indeterminate start and end points in mature muscle fibers (19,22). These MT lattices are mainly located in the cortex of muscle fibers, forming a supportive framework that preserves the internal structure of the muscle fiber (7,18,19). Such distinct reorganization of the MT network reflects not only developmental progression but also mechanical adaptation, making MT organization a valuable indicator of muscle fiber maturity and health. MTs have recently emerged as important regulators of muscle fiber mechanics (9,18,19,22). MTs are dynamic filaments whose stability and mechanical properties are modulated by post-translational modifications (PTMs) (18,23). For instance, acetylation of α-tubulin (acetyl-MT) at lysine-40 enhances MT resilience to contraction induced mechanical stress by increasing MT stiffness and consequently slowing down sarcomeric contraction and relaxation (10,24,25). In contrast, tyrosinated α-tubulin (Tyr-MT) marks dynamic or growing MTs, whereas detyrosinated α-tubulin (deTyr-MT) indicates stable, long-lived filaments (24,26). Functionally, deTyr-MTs promote desmin-mediated anchoring of the MT network to the sarcomere, forming load-bearing arrays that slow contraction and relaxation (11,27,28). This dynamic nature underlies MT’s involvement in diverse functions in muscle fibers, including maintenance of cell– and organelle-morphology, organelle positioning, mechanotransduction, and acting as a transport route for vesicles, organelles and mRNA to specific sites in the muscle fiber (5,22,29). Yet, the question remains how MT organization and its post-translational modifications (PTMs) quantitatively shape contractile output.

In our previous work, we established an *ex vivo* muscle fiber culture system that preserves long-term muscle fiber contractile activity by embedding cells in fibrin-based hydrogels, thereby limiting dedifferentiation (30,31). Notably, even where contractility was maintained, we observed progressive remodeling of MT architecture, largely by a loss in the lattice structure, suggesting that the organization of the MT network, rather than simply abundance, determines muscle fiber contractility. Because fibers in this system are removed from experiencing normal mechanical load, this provides a unique model to determine the MT remodeling that accompanies reduced contractile demand, a process relevant for disuse atrophy and aging (32–34). Here, we leverage this platform to dissect the relationship between MT organization, MT-PTMs, and muscle fiber function in a non-diseased context. We identify MT organization as a key structural determinant of sarcomere contractility and show that contractile decline follows a non-linear relationship with MT lattice disruption, independent of MT abundance. We further demonstrate that 3D culture conditions bias MT remodeling toward alignment along the long axis of the muscle fiber and reduce the fraction of deTyr-MTs, thereby alleviating internal mechanical constraints to preserve fiber contractility. By identifying deTyr– MT as a tunable regulator of this balance, our findings have potential relevance for approaches to counteract the loss of contractile function observed in aging and disuse atrophy.

## Methods

### Muscle fiber isolation

Post-mortem skeletal muscle tissue was obtained from animals sacrificed as part of other ethically approved research projects or breeding surpluses at the VU University in compliance with the European Council Directive (2010/63/ EU) by permission of the Animal Research Law of the Netherlands. Mature single muscle fibers were isolated following the protocol described by Vonk et al. (2023) (31). Briefly, flexor digitorum brevis (FDB) muscles were dissected from adult wild-type C57BL/6 mice (age 3-5 months) obtained from the Amsterdam Animal Research Centre and transferred to pre-warmed (37°C) and pH equilibrated (5% CO_2_) dissection medium, consisting of Minimum Essential Medium (MEM) with high glucose and pyruvate (Gibco, Thermo Fisher), 10% heat-inactivated fetal bovine serum (FBS, Thermo Fisher) and 1% penicillin-streptomycin (Sigma Aldrich). Muscles were digested in 0.2% w/v collagenase type II (Worthington Biochemicals) for 70-80 min at 37°C and 5% CO_2_, followed by incubation in 3ml of fresh dissection medium for 30 min. Individual muscle fibers were released by gentle trituration using two p1000 pipette tips with progressively narrower openings, and residual mononucleated cells and tissue were removed by gravity sedimentation.

### Muscle fiber culture

Isolated FDB fibers were maintained in a culture medium composed of MEM with high glucose and pyruvate (Gibco, Thermo Fisher) supplemented with 0.4% w/v Serum Replacement 2 (50X) (Sigma Aldrich), 1% w/v Horse Serum (Sigma Aldrich), and 1% w/v penicillin/streptomycin (Sigma Aldrich) and maintained at 37°C and 5% CO_2_. For 2D culture, polymer coverslip plates (Ibidi) were coated with mouse laminin (1:25 in MEM, Sigma Merck) for 2h before seeding. Half-media changes were performed on days 1, 3, and 5 post-isolation unless otherwise indicated.

### Muscle fiber embedding in hydrogel preparation and seeding

For 3D culture, isolated muscle fibers were embedded immediately after isolation in either increasing concentrations of Matrigel GFR Membrane Matrix (Matrigel, Corning) or a fibrin-based hydrogel. For Matrigel cultures, a muscle fiber pellet was mixed with Matrigel at final concentrations of 2mg/ml (denoted as “Low-laminin”) or 4mg/ml (denoted as “High-laminin”) and cast in culture dishes. Gels were prepared to cover 132 μL gel/cm^2^. For fibrin-based cultures, muscle fibers were embedded as described by Vonk et al. (2025) (30). Briefly, a two-step mixing procedure was used to prepare a final matrix composition of 2.5mg/mL fibrin and 10% of 4mg/mL Geltrex LDEV-Free Reduced Growth Factor Basement Membrane Matrix (Geltrex, Thermo Fisher) diluted in Iscove’s Modified Dulbecco’s Medium + L-glutamine (IMDM, Thermo Fisher). A “Gel solution” containing culture medium supplemented with 20% of 4mg/mL Geltrex, and 5.0mg/ml fibrinogen was prepared separately from a “Crosslinking solution” containing culture medium supplemented with 60μM aprotinin (a fibrinolysis inhibitor to prevent hydrogel degradation) (Thermo Fisher) and 0.5U/ml thrombin (Thermo Fisher). Both solutions were kept on ice until use. Muscle fiber pellets were resuspended in “Crosslinker solution”. Equal volumes of gel solution and crosslinking solutions were mixed briefly and transferred to culture dishes. Both Matrigel and fibrin gels were allowed to solidify for up to 2h at 37°C and 5% CO_2_, after which they were fully covered with culture medium. For fibrin cultures, the medium was supplemented with 60μM aprotinin and 200μM tranexamic acid (Thermo Fisher) to prevent degradation of the fibrin gel.

### Microtubule network manipulation

To alter the microtubule network, muscle fibers were treated with 100nM Taxol (Cayman Chemical) overnight to stabilize, 10µM parthenolide (PTL, Sigma Aldrich) for 2h to inhibit detyrosination or DMSO vehicle control. Changes in the MT network were verified by immunofluorescent staining for α-tubulin, detyrosinated-tubulin, and acetylated-tubulin.

### Electrical stimulation and contractile measurements

Muscle fibers kept in culture media were electrically stimulated at room temperature (RT) either at 2 hours (Day 0), 1, 3, 5 or 7 days post-isolation using a 6-well C-Dish insert (IonOptix) fitted into Corning 24-well plates and connected to a Myopacer cell stimulator (IonOptix). Stimulation was delivered as bipolar pulses at 10V, 1.0Hz frequency and a pulse duration of 4.0ms. Contractile properties were measured using the CytoCypher MultiCell High-Throughput System (IonOptix) (35). Sarcomere length was measured during contractions at a sampling rate of 350Hz. From these measurements, a contraction transient is generated, recording parameters such as sarcomere length, contractile velocity and contraction duration. For each muscle fiber, measurements were averaged across 9-10 consecutive contractions. Automatic data analysis was performed using the CytoSolver Transient Analysis Tool software. Transients with poor curve fitting (R^2^ < 0.95) were excluded from further analysis. For measurements where transients were rejected by the software, a cut-off of at least 4 transients was maintained, and data points whose average was based on 3 or fewer transients was removed from the dataset. The following contractile parameters were quantified: sarcomere length at rest, contractile velocity, contraction duration, % of sarcomere shortening, relaxation velocity, relaxation duration, and sarcomere length at peak contraction. Fractional sarcomere shortening was calculated on a per-fiber basis as the difference between resting and peak sarcomere length normalized to resting length, such that this paired, within-fiber metric is insensitive to the between-fiber variability in absolute sarcomere length that affects unpaired comparisons of resting or peak length across conditions.

### Immunofluorescent and live-cell labeling

Muscle fibers were fixed by replacing half of the culture medium with 4% paraformaldehyde (PFA, Thermo Fischer) for 5 min, followed by fresh 4% PFA for 10 min. Samples were washed three times for 5 min with PBS and stored at 4°C for up to 7 days. For blocking and permeabilization, fibers were incubated for 1h at 4°C on a shaker in blocking buffer containing 3% bovine serum albumin (BSA, Sigma Aldrich), 0.2% Triton-X100 (Sigma Aldrich) and 0.05% Tween (Sigma Aldrich) dissolved in PBS. Samples were incubated in blocking buffer with added primary antibodies (diluted according to manufacturer’s instructions, **Table 1**) overnight at 4°C on a shaker. After washing three times for 5 minutes with IF buffer containing 0.3% BSA, 0.2% Triton-X100 and 0.05% Tween in PBS, samples were incubated with AlexaFluor secondary antibodies (1:250, Invitrogen), DAPI (1:1000, Invitrogen) and AlexaFluor Plus 647 Phalloidin (1:1000, Invitrogen) for 1h. Excess staining solution was removed by washing the samples three times for 5 min with IF buffer, and samples were stored in PBS at 4°C for up to 5 days. For live cell imaging of MTs, muscle fibers were incubated with SPY650-tubulin (1:1000, Spirochrome) for 2h and washed with phenol red-free IMDM (ThermoFischer) and imaged within 2h at RT.

**Table 1:** Antibodies used for immunofluorescent labeling.

| Antibody reference name | Antigen | Antibody isotype | Dilution | Company |
| --- | --- | --- | --- | --- |
| 12G10 | α-tubulin | Mouse IgG1 | 1μg/mL | Developmental Systems Hybridoma Bank |
| T7451 | Acetylated-tubulin | Mouse IgG2b | 1:200 | Sigma Aldrich |
| ab48389 | Detyrosinated-tubulin | Rabbit | 1:200 | Abcam |

### Fixed cell image acquisition

Images were acquired using confocal microscope (Nikon Ti2 ZDrive) equipped with a Prime BSI camera (A21A726014), CrestOptics X-Light spinning disc system with a 50µm pinhole (CrEST X-Light V3), laserbox (NIDAQ AOTF Multilaser) with line wavelengths of 405nm, 470nm, 555nm and 640nm. The muscle fibers were imaged at a magnification of 10X (plan Fluor 10X OFN25 DIC N1) or 60X (PLAN APO 60X IOL OFN25 DIC). Z-stacks were collected at 1μm intervals and processed in ImageJ to generate maximum-intensity projection images.

### Image quantification

Maximum Intensity projections (MIP) were generated using ImageJ unless otherwise indicated. Phalloidin staining was used to define muscle fiber boundaries by generating a mask and applying a homomorphic filter to correct for non-uniform illumination. Viability was measured by counting the number of viable muscle fibers in a 24-well plate on Days 1 and 7. The viable cell count as a percentage was determined by dividing the cell count on Day 7 by Day 1. An overview image of a single 24-well was obtained by stitching MIP images with 10% overlap acquired with a 10X objective. Mean fluorescence intensity was quantified within each isolated muscle fiber by generating a mask (Otsu method) and measuring the intensity within the masked area. To obtain the area of each individual fiber, we used the ImageJ option analyze particle with the additional selection criteria of size selection (10 cm^2^-infinity), include holes, and add to manager to save each mask as a new region of interest. Grey values were determined in ImageJ, using the set measurement options area and mean grey value. For ratio measurements, the mean grey values within the masked regions were divided by each other. The MT-network was classified using a MATLAB script written for blinded classification quantification. The MT networks within cultured single muscle fibers were randomized and presented to 5 participants, blinded to the experimental conditions. Participants were asked to classify MT-lattice organization to the predetermined groups: A) organized, B) semi-organized or C) random. The majority vote out of all 5 participants was used to assign the corresponding group to classify the MT network organization. MATLAB script for blinded classification will be shared upon request. MT organization was quantified using the Texture Directionality Tool (TeDT) developed by Liu and Ralston (2014) (21). From the angular histogram, the longitudinal and transverse MT oriented population was extrapolated. The longitudinal MT population was computed by averaging the values from bins (0°, 4°, 172° and 176°) and the transverse MT population was obtained by averaging the values from bins (84°, 88°, 92°, 96°).

### Correlation microtubule network to contractility

Muscle fibers were incubated with SPY650-tubulin (1:1000, SpyroChrome) overnight and washed with phenol red-free IMDM (Thermo Fisher) and imaged within 2h at room at RT on Day 3. Muscle fibers were electrically stimulated using a 35 mm Dish insert (IonOptix) fitted into a 35 mm high polymer coverslip dish (Ibidi) and connected to a Myopacer cell stimulator (IonOptix). Stimulation was delivered as bipolar pulses at 10V, 1.0Hz frequency and a pulse duration of 4.0ms.

Muscle fiber sarcomere striation was visualized using brightfield imaging at a sampling rate of 400 frames per second for 10 seconds. After obtaining the electrical stimulation-induced sarcomere recording, the electrical stimulation was stopped and the MT network was imaged. Sarcomere shortening was measured using the ImageJ tool SarcOptiM (High Frequency Online Sarcomere Length Measurement), MT abundance was measured by the mean intensity, and the MT-lattice was measured using the TeDT method. All model fittings were performed using GraphPad Prism’s XY analyses tool for simple linear regression and nonlinear regression (curve fit) and confirmed in MATLAB fitting the obtained equation from Prism on to the dataset.

### Statistical analysis

Experimental results from muscle fibers were obtained from at least three separate isolations and 3 individual mice, unless otherwise stated. For single fiber experiments, n is denoted as the number of fibers, while N is denoted as the number of mice. Mean values were generated for each mouse, and statistical analysis was performed using either a linear mixed model or ordinary one-way *ANOVA* with a Tukey post-hoc test. For those tested with a linear mixed model each animal was defined as a random intercept to correct for multiple measurements taken from the same animal and a Tukey post-hoc test was used to correct for multiple comparison. A threshold p-value of *p* < 0.05 was considered significant. All statistical analyses were performed using RStudio.

## Results

### MT organization and sarcomere contractility progressively decline over 7 days of 2D ex vivo culture

We previously showed that 7 days of *ex vivo* culture in 2D disrupts the MT lattice and reduces sarcomere shortening by ∼50% (30). To get a better understanding of the temporal changes in MT organization over time, isolated muscle fibers were cultured in 2D for up to 7 days, fixed at various time points, and immunolabeled for α-tubulin (**Figure 1A**). A clear MT lattice was evident on Days 0 and 1, but gradually diminished thereafter and was largely absent by Day 5. Blinded classification into organized, semi-organized and random MT patterns confirmed this trajectory (**Figure 1B**), with more than 60% of muscle fibers being classified as organized at Day 0, which then declined in an almost linear fashion with culture duration, leading to almost no muscle fibers having an organized MT network at Day 7. Semi-organized muscle fibers persisted throughout, accounting for 20-50% of the population, with near-equal representation of all classes observed on Day 3.

**Figure 1:**
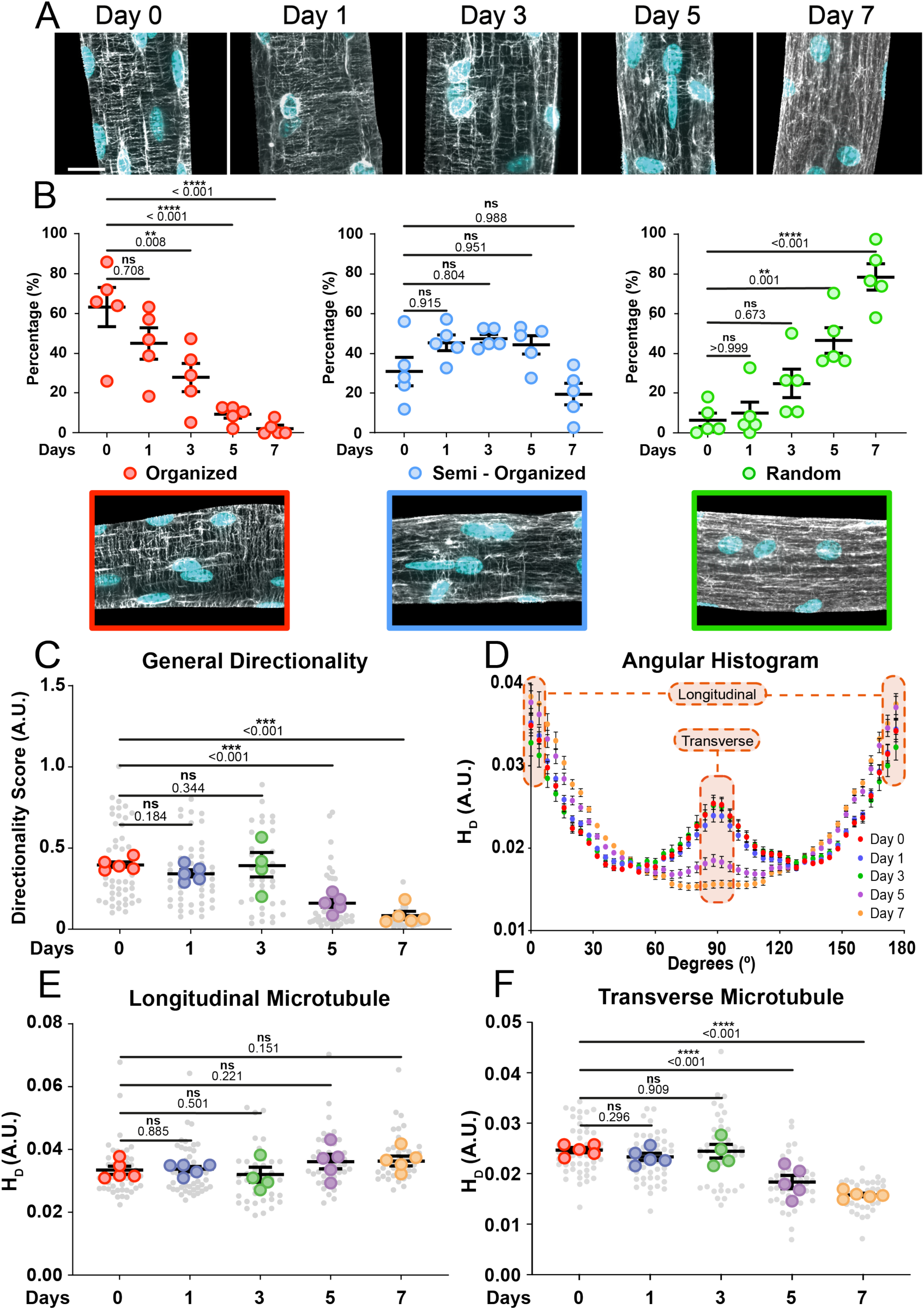
Muscle fiber microtubule organization becomes disrupted during *ex vivo* culture, primarily through the progressive loss of transverse microtubules: **(A)** Representative maximum intensity projections of muscle fiber at various time points during *ex vivo* culture immunofluorescently labeled for α-tubulin (grey) and nuclei (cyan). Scale bar = 20μm **(B)** Blinded classification of microtubule organization of muscle fibers cultured up to 7 days *ex vivo*. **(C)** General directionality score of microtubule organization within fibers cultured up to 7 days *ex vivo*. **(D)** Angular histogram of the microtubule direction within muscle fibers cultured up to 7 days *ex vivo*. **(E,F)** From the angular histogram, extrapolated data to quantify the longitudinal and transverse microtubule organization within muscle fibers cultured up to 7 days *ex vivo*. Data are means ± SEM; *N* = 5 participants in panel B and *N* = 5 mice in panel C-F with *n* = muscle fibers. Significance was determined using a one-way ANOVA for panel B and linear-mixed model for panels C, E and F with *p* < 0.05 considered significant with * = *p* < 0.05, ** = *p* < 0.01, *** = *p* < 0.001, and **** = *p* < 0.0001.

To assess whether MT remodeling followed a defined pattern, we applied the Texture Directionality Tool (TeDT) method developed by Liu and Ralston (2014) (21). This tool measures filament directionality as a global descriptor of lattice organization, while also allowing for the extraction of MTs with a specific orientation. Using this approach, we found that the general directionality score remained stable from Day 0 to 3, indicating preservation of the MT lattice (**Figure 1A,C**). On Day 5, there was a marked decrease in the organization of the MT lattice, with the directionality score decreasing by ∼60%, (**Figure 1C**), and on Day 7, the directionality score was close to 0.00, consistent with a highly disorganized MT network (**Figure 1A,C**). To determine whether MT lattice loss reflected a redistribution of specific MT orientations, we extrapolated from angular histograms the longitudinal and transverse MT populations. The angular histograms showed the expected peaks for longitudinal (0° and 180°) and transverse (90°) MTs (**Figure 1D**) (13,21). Longitudinal MTs remained stable across the 7 days (**Figure 1E**), whereas transverse MT decreased by ∼25% on Day 5 and ∼35% on Day 7 (**Figure 1F**), paralleling the general directionality score decline (**Figure 1C**). These findings suggest the loss of transverse MTs drives the global disruption of MT lattice during *ex vivo* culture, with missing transverse elements redistributed across a broad range of angles that yield the disorganized MT network observed in long-term 2D culture (**Figure 1A**).

To determine whether there were changes in the MT network abundance, we quantified α-tubulin mean fluorescence intensities at each time point. Total MT network abundance increased by ∼40% during the first 3 days, followed by a ∼20% reduction between Days 3 and 5, and subsequent ∼15% increase by Day 7, nearly reaching Day 3 levels (**Figure S1**). The divergence between general directionality and MT network abundance in time indicates that MT network remodeling is independent of MT filament abundance.

### Changes in sarcomere contractility follow the same temporal decline as MT organization

To determine whether the observed changes in MT organization were associated with changes in muscle fiber contractile function, sarcomere shortening and kinetics were measured in unloaded muscle fibers using our established electrical stimulation protocol (**Figure 2A**) (31). Sarcomere shortening declined by ∼50%, from ∼10% on Day 0 to ∼5% on Day 7 (**Figure 2B,C & Figure S2C**), corroborating previous findings (30). Although non-significant, a modest (∼10%) increase in shortening was observed between Day 0 and Day 1, possibly reflecting fiber recovery after enzymatic isolation. Interestingly, sarcomere shortening declined linearly (R^2^ = 0.81, **Figure S3**) between Day 1 and Day 7. The decrease in shortening was primarily driven by the change in resting sarcomere length, which decreased by ∼5% from 1.92μm on Day 0 to 1.83μm on Day 7, plateauing by Day 5 (**Figure 2B & Figure S2A**). Sarcomere length at peak contraction remained largely unchanged throughout the culture period, averaging 1.73-1.77μm (**Figure 2B & Figure S2B**). Relaxation velocity was preserved through Day 3 but declined rapidly by ∼90% by Day 7, decreasing from 4.9μm/s on Day 3 to 0.2μm/s on Day 7 (**Figure 2D**). This was accompanied by a ∼50% prolongation in relaxation time to reach 70% baseline sarcomere (**Figure 2E**).

**Figure 2:**
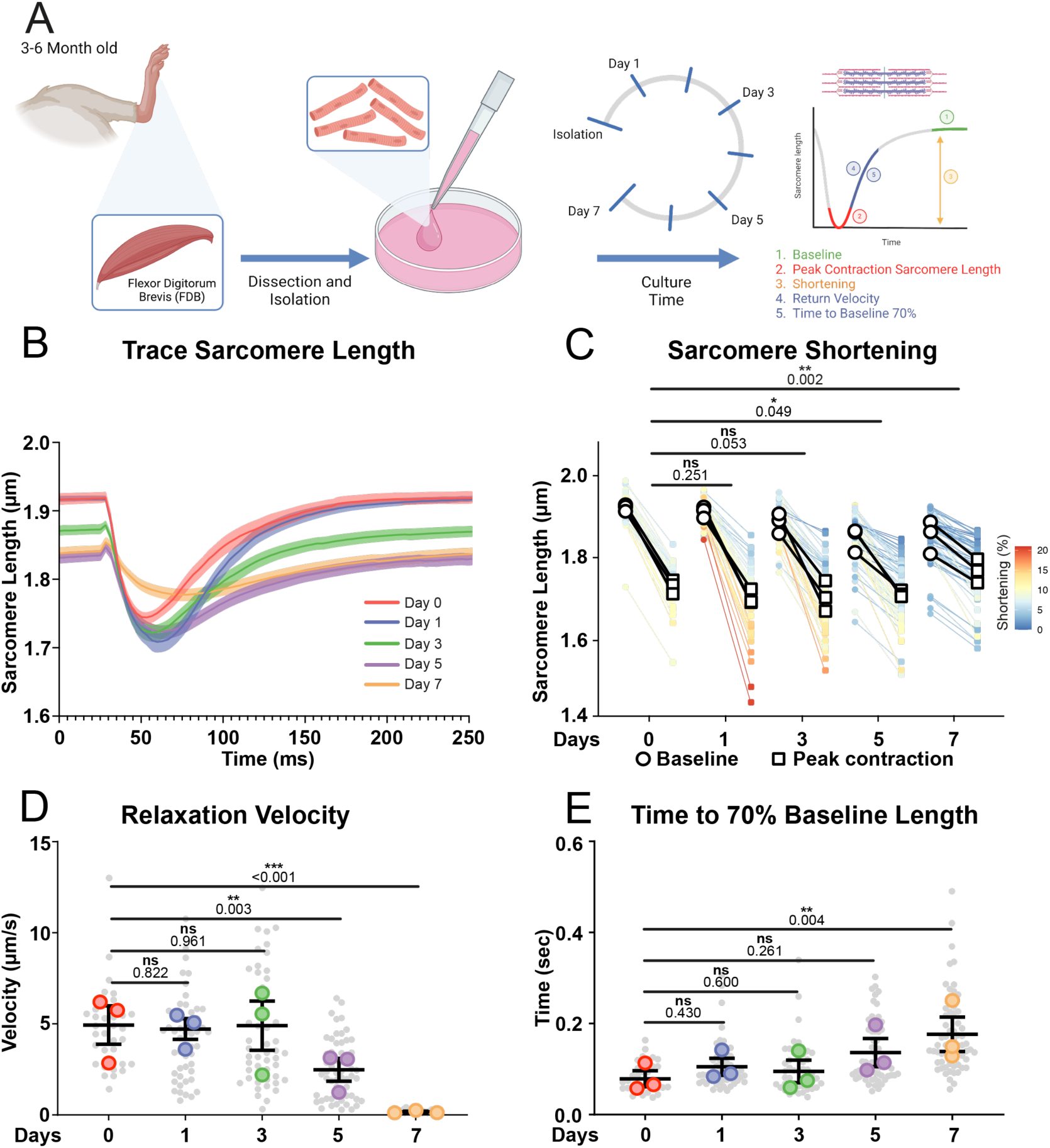
Progressive loss of sarcomere contraction during 2D *ex vivo* cultured muscle fibers. (A) Graphical overview of the culture experiment (Image made in Biorender). **(B)** Traces of sarcomere length measured during electrically-stimulated sarcomere contraction within intact muscle fibers kept in 2D culture on laminin-coated dishes up to 7 days *ex vivo*. **(C)** Contractile measurements of muscle fibers kept in culture on laminin-coated dishes up to 7 days *ex vivo*. The resting sarcomere length (baseline) and sarcomere length during maximal contraction (peak sarcomere) were measured and used to compute the percentage of shortening depicted in the heat-map. The statistics are run on the percentage shortening. Individual plots for baseline sarcomere length, sarcomere length at peak contraction, and percentage shortening are depicted in Figure S2. **(D)** Quantification of the sarcomere relaxation at increasing culture time duration. **(E)** Quantification of relaxation time to 70% of the baseline of fibers cultured up to 7 days *ex vivo*. Data are means ± SEM; *N* = 3 mice and *n* = fiber. Significance was determined using a linear-mixed model with *p* < 0.05 considered significant with * = *p* < 0.05, ** = *p* < 0.01, *** = *p* < 0.001, and **** = *p* < 0.0001.

Collectively, these data show that isolated muscle fibers retain their MT lattice and sarcomere contractility for up to 3 days *ex vivo*, after which progressive MT remodeling and contractility decline occur under 2D conditions, largely due to changes in resting sarcomere length. The loss of contractile amplitude follows a near-linear trend over time, while relaxation kinetics deteriorate sharply once contractile performance becomes compromised. The temporal overlap between MT remodeling and contractile dysfunction suggests a relationship between MT organization and sarcomere contractility.

### Microtubule organization predicts contractile performance in isolated muscle fibers

To investigate whether MT organization is associated with contractile performance within individual *ex vivo* cultured muscle fibers, we analyzed both parameters in the same muscle fiber at Day 3 of 2D culture. This time point was selected to capture the pronounced heterogeneity in both MT organization (**Figure 1**) and contractility (**Figure 2**). We measured sarcomere shortening via striation spacing during electrical stimulation and visualized MTs live using a SPY-tubulin probe (**Figure 3A & Figure S4A,B**). Because SPY-tubulin uses a Taxol-based labeling approach, and Taxol is known to influence MT organization (11,30), we first determined whether the Taxol component within the live cell-dye we were using altered MT organization in muscle fiber by measuring the MT directionality. Directionality scores obtained from SPY-tubulin-labeled fibers (**Figure S4C,D**) closely matched those of antibody-labeled fixed muscle fibers (**Figure 1C,D**), indicating negligible influence from the dye. SPY-tubulin probes are smaller and more cell-permeable than conventional immunofluorescent antibodies, enabling faster penetration into cells and visualization of both the cortical and core MT network (**Figure S4A**) (5). This dual-domain imaging complicates MT organization assessment. To address this, imaging was restricted to a single focal plane focusing on the MT core, which is less densely packed and exhibits more apparent MT grid-like structures (**Figure S4B**). Slightly higher scores observed in SPY-tubulin images likely reflect improved filament resolution as separation of individual MT filaments in the loosely packed MT core (**Figure S4B**) is much higher when compared to the dense MT cortex (**Figure 1A & Figure S4A**) (5).

**Figure 3:**
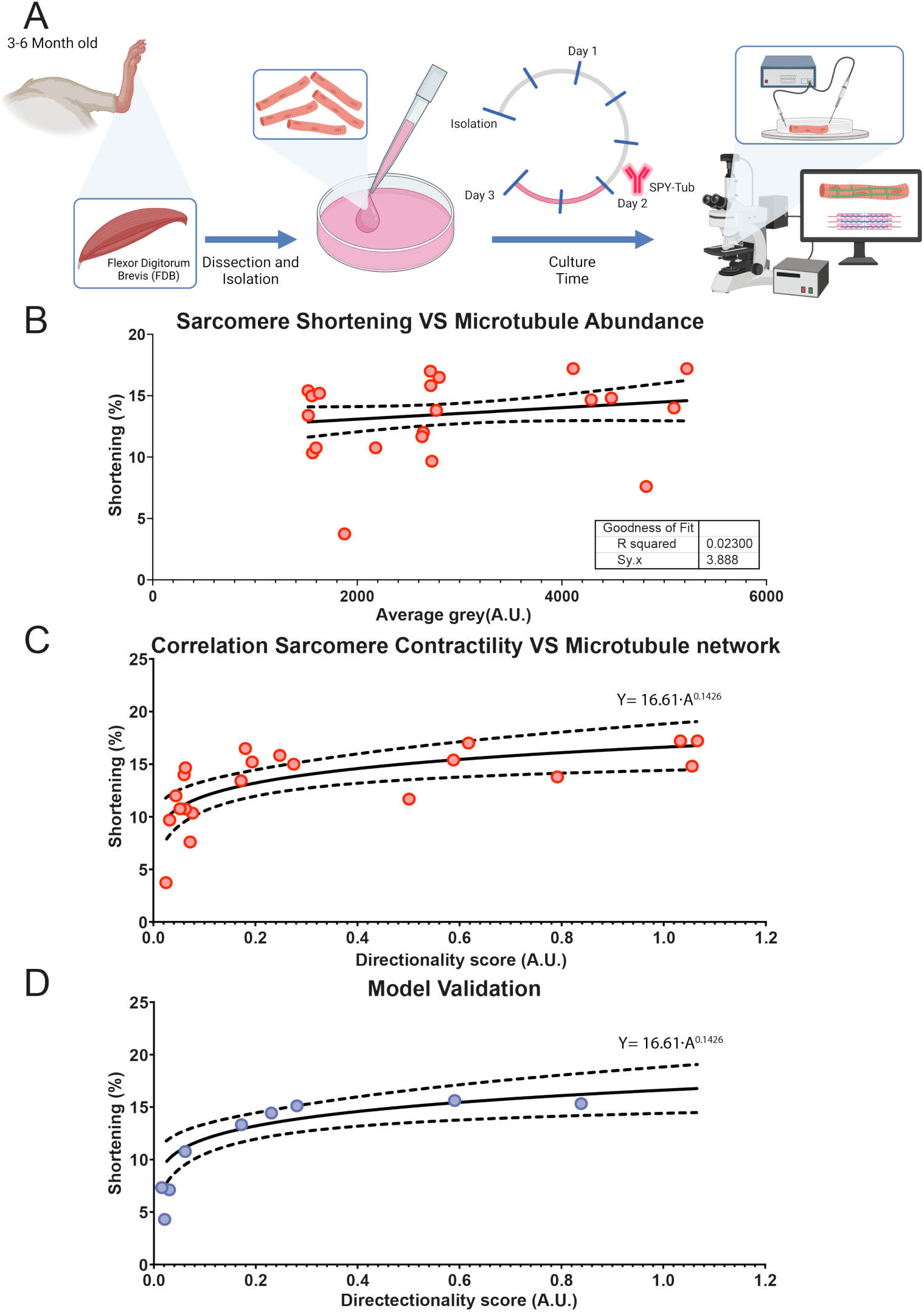
Sarcomere shortening of *ex vivo* cultured muscle fibers can be predicted from microtubule organization. **(A)** Graphical overview of the culture experiment to correlate sarcomere shortening with microtubules (Image made in Biorender). **(B)** Linear correlation analysis between sarcomere shortening and microtubule abundance, defined as average grey value, in muscle fibers kept in culture for 3 days *ex vivo*. **(C)** Non-linear correlation between sarcomere shortening and microtubule organization, defined as a general directionality score, in muscle fibers kept in culture for 3 days *ex vivo*. **(D)** Validation of the predicted correlation between sarcomere shortening and microtubule organization with a new data set.

After confirming that SPY-tubulin did not alter MT organization, we analyzed contractility, total MT filament abundance and core MT organization within single muscle fibers. Linear regression revealed no correlation between contractility and MT abundance (R^2^ = 0.02, **Figure 3B**). In contrast, the MT organization showed a non-linear relationship with contractility (**Figure 3C**). A power-law model best described the data and identified a critical transition near a directionality score of ∼0.1. Muscle fibers with a score above this threshold retained sarcomere shortening, whereas those below ∼0.1 exhibited marked MT disorganization and reduced sarcomere shortening. The model’s predictive robustness was confirmed with an independent dataset, which closely overlapped the initial curve (**Figure 3D**), validating MT organization as a reliable predictor of sarcomere contractility under 2D culture conditions. Together, these findings demonstrate that MT organization, rather than total MT abundance, predicts contractile performance in *ex vivo* cultured muscle fibers. The data further identify a directionality threshold of ∼0.1 that distinguishes muscle fibers with preserved contractile function from those with compromised contractile function.

### 3D culture preserves contractility while shifting MTs toward longitudinal alignment

We previously reported that culturing muscle fibers in a fibrin-based hydrogel (3D) leads to preserved contractile function during *ex vivo* culture (30). Having established a link between MT organization and muscle fiber contractility (**Figure 3**), we next investigated whether 3D culture could preserve contractility in part by preventing MT remodeling. Focal adhesions and laminin-mediated membrane attachments anchor MTs via adaptor proteins to the sarcolemma (9,16,18). Their disruption in disease states compromises MT anchoring, MT lattice disorganization, and increased MT density (9,16,18), similar to what we are observing with prolonged *ex vivo* culture. We therefore tested whether increasing laminin-rich matrix may increase laminin-mediated ECM interactions to stabilize MT organization during prolonged *ex vivo* culture. To do this, we used basement membrane extracts (BME) derived from Engelbreth-Holm-Swarm (EHS) tumors (Matrigel and Geltrex) to modulate the laminin content within gels, as laminin constitutes more than half of their protein composition (36). Isolated muscle fibers were cultured in laminin low (2 mg/mL Matrigel) and laminin high (4.0 mg/mL Matrigel) hydrogels, fibrin-based hydrogels (containing 0.5 mg/mL Geltrex), and compared with 2D as a reference condition (30,31). Muscle fiber viability after 7 days was significantly improved in all gel conditions (∼85% viability) compared to 2D culture (∼50% viability) (**Figure S5**), consistent with our previous report (30). In the absence of laminin in the gel composition, muscle fibers hypercontracted within 24 hours, indicating that a laminin-containing matrix is required for sustained viability in 3D cultures.

We next analyzed MT organization across conditions. General directionality scores followed a similar trend across all conditions (**Figure 4A, & Figure S6A,B**), indicating preservation of the MT lattice up to Day 3 independent of culture condition, followed by a significant decrease in organization at Day 5. These data suggest that increasing laminin-mediated ECM interactions through higher laminin content in gels was not sufficient to preserve MT lattice integrity. However, analysis of the angular histogram revealed a shift in the MT orientation in muscle fibers cultured in 3D (**Figure S7A-D**). When MT orientation was further resolved into the longitudinal vs transverse population at Day 1 and Day 7 (**Figure 4B,C & Figure S7E,F**), the transverse MT population was markedly reduced in all conditions at Day 7. Relative to 2D culture, 3D cultures showed an increase in their longitudinal MT population by ∼35% (**Figure 4B & Figure S7E**), while the transverse MT population dropped by ∼30% (**Figure 4C & Figure S7F**) on Day 7. These measurements indicate an axial bias towards longitudinal MT alignment in 3D culture, independent of laminin dose, rather than isotropic dispersion of MTs observed in 2D during prolonged *ex vivo* culture of muscle fibers.

**Figure 4:**
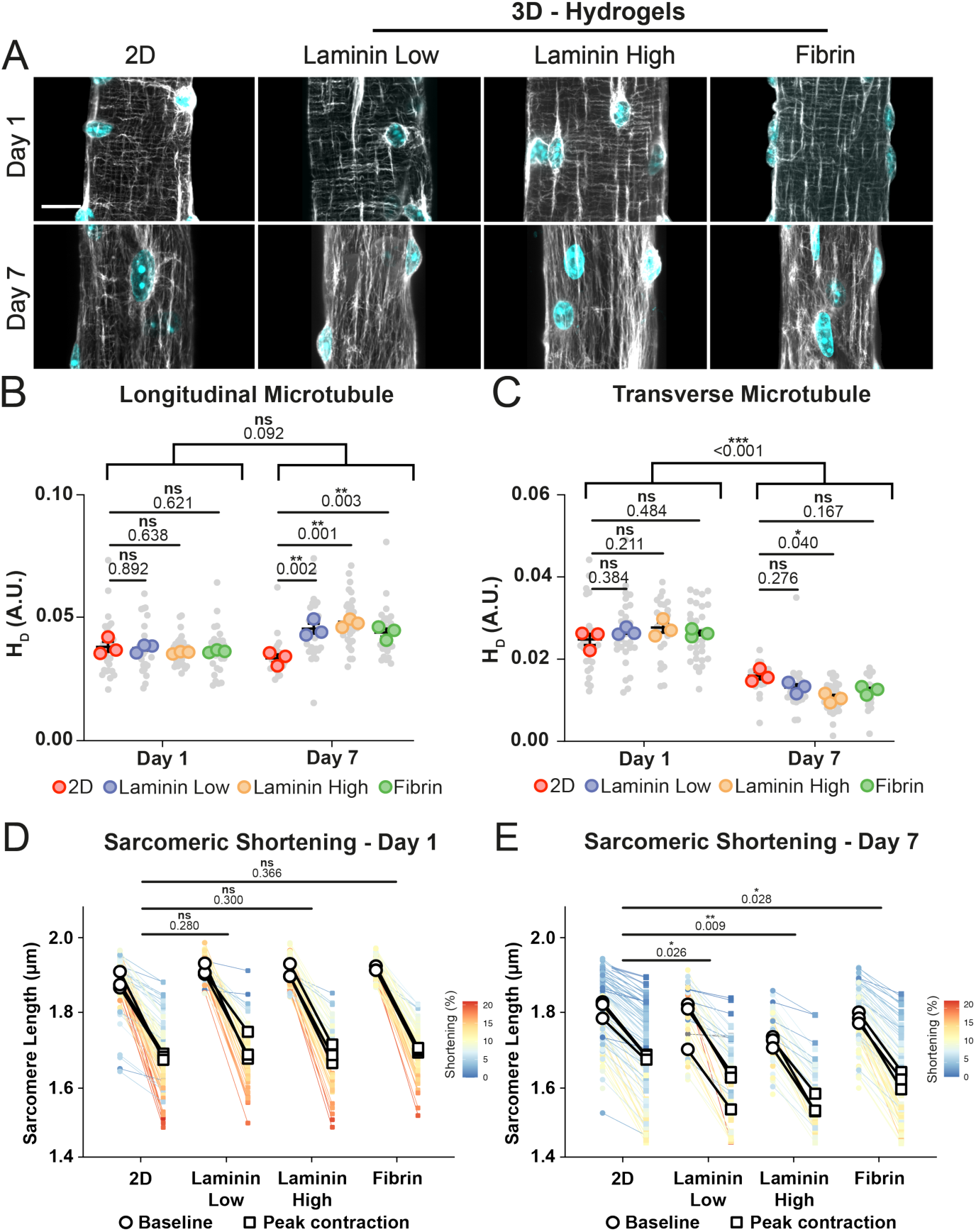
3D culture leads to remodeling of the MT lattice that favors longitudinal MT orientation, and prevents the time-dependent decline in sarcomeric shortening, independent of the laminin concentration. **(A)** Representative maximum intensity projections of muscle fibers at Day 1 and Day 7 during *ex vivo* culture immunofluorescently labeled for a-tubulin (grey) and nuclei (cyan). Scale bar = 20μm **(B)** From the angular histogram extrapolated data to quantify the longitudinal microtubule organization within muscle fibers on Day 1 and Day 7 **(C)** From the angular histogram extrapolated data to quantify the transverse microtubule organization within muscle fibers on Day 1 and Day 7. **(D)** Contractile measurements of muscle fibers on Day 1. **(E)** Contractile measurements of muscle fibers on Day 7. The resting sarcomere length (baseline) and sarcomere length during maximal contraction (peak sarcomere) were measured and used to compute the percentage of shortening depicted in the head-map. The statistics are run on the percentage shortening. Individual plots for baseline sarcomere length, sarcomere length at peak contraction, and percentage shortening are depicted in Figure S8. Data are means ± SEM; *N* = 3 mice with *n* = muscle fibers. Significance was determined using a linear-mixed model with *p* < 0.05 considered significant with * = *p* < 0.05, ** = *p* < 0.01, *** = *p* < 0.001, and **** = *p* < 0.0001.

In parallel, we quantified contractile parameters in 2D and 3D cultures on Days 1 and 7. In line with our previous report (30), in all culture conditions, resting sarcomere length, sarcomere length at peak contraction, and sarcomere shortening were equivalent on Day 1, averaging ∼1.9μm, ∼1.7μm and ∼11%, respectively (**Figure 4D & Figure S8A,C,E**). On Day 7, resting sarcomere length decreased across all culture conditions (**Figure 4E & Figure S8B**), with the largest reduction observed in laminin high gels (∼10%), and lesser reductions in laminin low gels (∼7.5%), fibrin-based hydrogel (∼6.5%), and 2D (∼4.0%), relative to their respective Day 1 values (Error! Reference source not found.**D & Figure S8A**). Sarcomere length at peak contraction declined in a similar manner in the 3D conditions (**Figure 4E & Figure S8D**), with the most pronounced decline in laminin high gels (∼8%), followed by laminin low (∼5.5%) and fibrin-based hydrogel (∼5%) compared with Day 1. Despite these changes in absolute sarcomere length (Error! Reference source not found.**E**), sarcomere shortening was preserved across all 3D conditions at Day 7 (Error! Reference source not found.**E & Figure S8F**), averaging ∼9.5%. This preservation arises because, unlike in 2D where peak contraction length remained fixed while resting length declined, sarcomere length at peak contraction in 3D fell in proportion to resting length. In contrast, 2D cultured muscle fibers exhibited a ∼40% loss of contractile amplitude (∼11% to 6%), as sarcomere length at peak contraction remained constant (∼1.7μm) while the resting sarcomere length dropped to ∼1.8μm over the same period, consistent with our previous report (30). Although, 3D culture preserved muscle fiber contractility, contractile kinetics (relaxation velocity) were altered over the 7 days culture period (**Figure S8G,H**). Despite being non-significant, embedding isolated muscle fibers in gels resulted in a ∼20% increase in relaxation velocity compared to 2D on Day 1 (**Figure S8G**). After 7 days of ex vivo culture, relaxation velocity decreased by ∼50% in all culture conditions, while the ∼20% difference between 2D and 3D was maintained (**Figure S8H**). These findings indicate that 3D matrices prevent the time-dependent decline in contractile performance observed under 2D conditions, and that contractility is independent of laminin concentration within the tested range. Together, these results demonstrate that laminin-containing 3D hydrogels maintain muscle fiber viability and contractility for at least 7 days *ex vivo*, thereby preventing the early functional deterioration characteristic of 2D culture and biasing MT orientation towards longitudinal alignment.

### Microtubules post-translational modifications reveal distinct stabilization patterns in prolonged culture

Since MT abundance and organization did not explain the increased contractile performance following 7 days of culture in 3D, we next assessed whether 2D and 3D culture conditions modify MT-PTMs, specifically acetylation and detyrosination. MTs enriched in acetyl-MT increase cytoskeletal stiffness and viscoelastic resistance in muscle fibers, leading to slower contraction kinetics and reduced fractional shortening (10). Similarly, deTyr-MT (removal of C-terminal tyrosine from α-tubulin) promotes desmin-mediated anchoring to the sarcomeres, thereby slowing contraction and relaxation in muscle fibers (11,27,28). To determine how the MT-PTM landscape evolves during prolonged *ex vivo* culture, we quantified acetyl-MT and deTyr-MT in 2D and 3D conditions, focusing on the fibrin hydrogel since neither MT organization (**Figure 4B,C**) nor contractility (**Figure 4D,E**) varied among gel formulations.

On Day 1, both 2D and 3D displayed a well-defined MT lattice with pronounced perinuclear enrichment of α-tubulin, acetyl-MT and deTyr-MT (**Figure 5A**). Visual inspection revealed that in the cytoplasm, deTyr-MTs were mainly localized along transverse filaments, whereas acetyl-MTs localization was more broadly distributed along both axes. Quantification of the total abundance showed similar α-tubulin, acetyl-MT and deTyr-MT levels between 2D and 3D fibers on Day 1 (**Figure 5A & Figure S9**). Measuring the ratio of acetyl-MT or deTyr-MT over α-tubulin was used as a fiber size-independent measure to gain insight into the MT-PTM fingerprint. Our measurements revealed no change in the acetyl-MT (**Figure 5A,B**) or deTyr-MT (**Figure 5A,C**) ratio on Day 1 in both culture conditions, indicating that 3D did not alter the initial MT-PTM profile.

**Figure 5:**
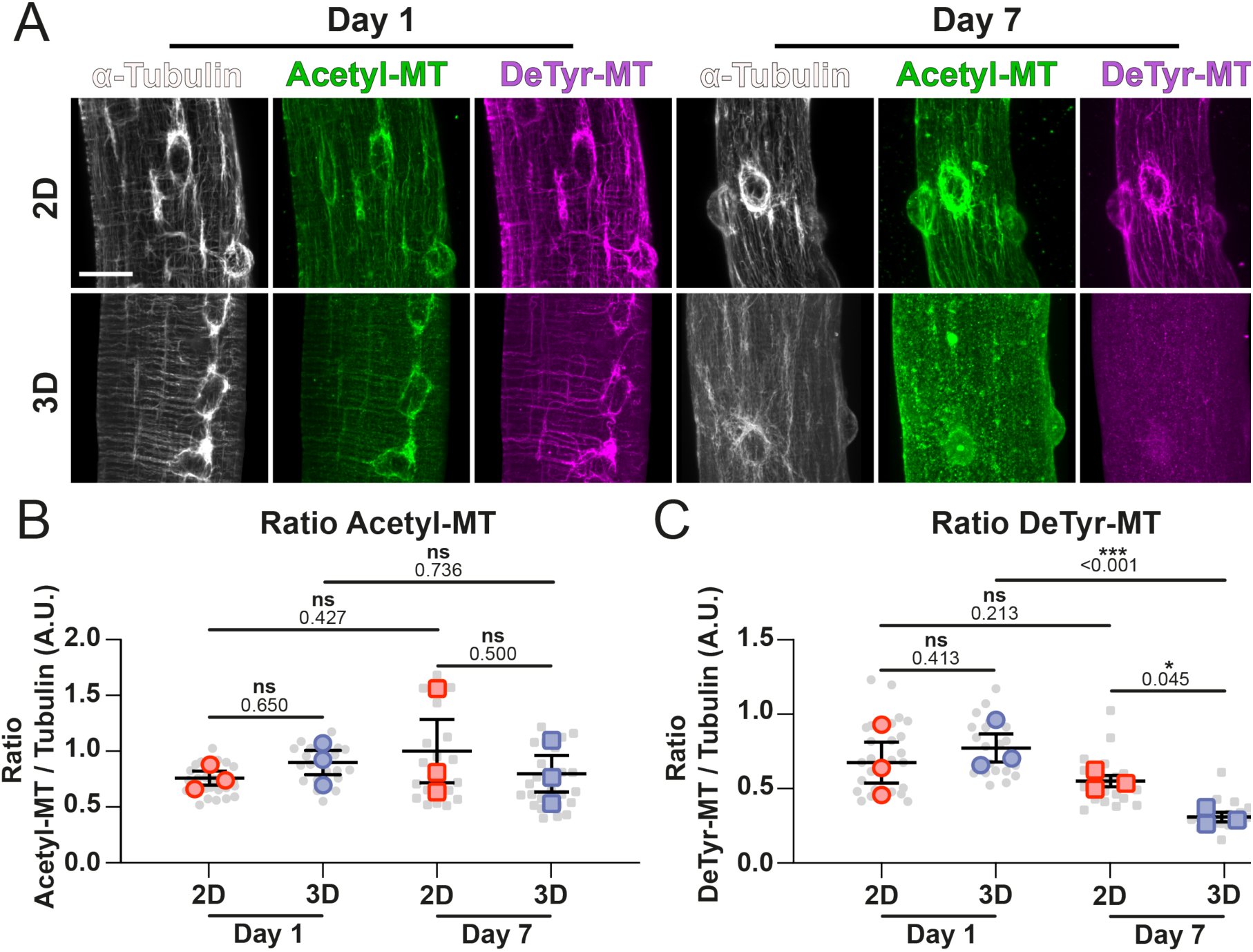
Long term culture in 3D declines the fraction of detyrosinated microtubules. **(A)** Representative maximum intensity projections of muscle fiber at either Day 1 or Day 7 immunofluorescently labeled for a-tubulin (grey), acetylated tubulin (Acetyl-MT) (green) and detyrosinated tubulin (DeTyr-MT) (cyan). Scale bar = 20μm **(B)** Quantification of the fluorescence intensity of the acetyl-MT fraction relative to a-tubulin in muscle fibers cultured in 2D and 3D on Days 1 and 7. **(C)** Quantification of the fluorescence intensity of the deTyr-MT fraction relative to a-tubulin in muscle fibers cultured in 2D and 3D on Days 1 and 7. Data are means ± SEM; *N* = 3 mice and *n* = fiber. Significance was determined using a linear-mixed model with *p* < 0.05 considered significant with * = *p* < 0.05 and *** = *p* < 0.001.

On Day 7, MT lattices had disassembled in both conditions in agreement with previous results (**Figure 4A & Figure 5A**). However, perinuclear enrichment of both MT-PTMs persisted only in 2D, whereas such perinuclear enrichment remained absent in 3D (**Figure 5A**). Total abundance increased over the 7 days *ex vivo* culture period in both conditions, but the increase was substantially greater in 3D (∼115%) than in 2D (∼35%) (**Figure S9A**), consistent with our earlier observations (**Figure S1**). This observation suggests that total MT content accumulates in time independent of the culture condition, yet the increase in MT abundance is more pronounced in 3D culture, thereby, 3D culture induces MT remodeling of the perinuclear region.

In the cytoplasm, acetyl-MTs remained broadly distributed in both conditions on Day 7, yet their localization patterns were different (**Figure 5A**). In 2D, acetyl-MTs remained present along the longitudinal MT filaments, while in 3D the staining showed a more dispersed acetyl-MT pattern lacking a clear organization. Despite the clear separation in acetyl-MT patterns between 2D and 3D cultures on Day 7, their ratio of acetyl-MT to total MT remained largely unchanged, near a ratio of 0.5 (**Figure 5B**). This suggests that acetyl-MT levels scale proportionally with the overall MT abundance, which was confirmed by measuring an increase in acetyl-MT abundance of ∼85% in 3D compared to ∼75% in 2D (**Figure S9B**). In contrast, deTyr-MT did not increase proportionally with MT abundance (**Figure S9C**), resulting in a progressive decline in the deTyr-MT fraction (**Figure 5C**) by Day 7. This decline was more pronounced in 3D cultures, as the fraction of deTyr-MT dropped by ∼50% to near undetectable levels on Day 7 (**Figure 5C & Figure S9C**). Together, these findings indicate that prolonged *ex vivo* culture selectively reshapes the MT-PTM landscape, and that embedding muscle fibers in a hydrogel appears to alter the dynamics of the MT network, resulting in a lower abundance of deTyr-MTs.

### Microtubule detyrosination levels are associated with improved contractility via two distinct routes

Having established that preserved contractility in 3D culture coincides with a reduced deTyr-MT fraction (**Figure 5**), we next sought to determine whether detyrosination actively governs contractile output rather than merely accompanying it. Addressing this would also clarify counterintuitive findings from our earlier work, where we found that pharmacologically stabilizing MTs with Taxol or destabilizing them with nocodazole produced similarly positive effects on muscle fiber contractility (30). Since MT abundance alone is not associated with function (**Figure 3B**), whereas both MT organization (**Figure 3**) and a PTM profile that reduces viscoelastic load (**Figure 5**) are, we reasoned that these opposing manipulations may improve contractility through separate mechanisms. That is, Taxol by reorganizing the MT lattice, and nocodazole by relieving viscoelastic resistance. To test the latter directly, we inhibited detyrosination with PTL rather than globally destabilizing the network, isolating the PTM contribution from changes in MT abundance. In parallel, muscle fibers were treated with Taxol to increase MT stability and promote detyrosination.

Immunofluorescence analysis for α-tubulin, acetyl-MT and deTyr-MT confirmed effective pharmacological modulation with PTL and Taxol in both 2D and 3D on Day 7 (**Figure 6A & Figure S10A-C**). Neither drug altered total MT abundance in either condition (**Figure S10A**), indicating that the treatment specifically affected MT-PTM state rather than MT abundance. As expected, PTL reduced deTyr-MT with a ∼30% decrease in 2D and a ∼10% decrease in 3D (**Figure S10B**), without affecting acetyl-MT (**Figure S10C**). This shifted the deTyr-MT fraction downward in 2D by ∼50%, bringing it close to the lower 3D baseline (**Figure 6B**). Consistent with this, PTL did not change the directionality score in either culture system (**Figure 6C**).

**Figure 6.**
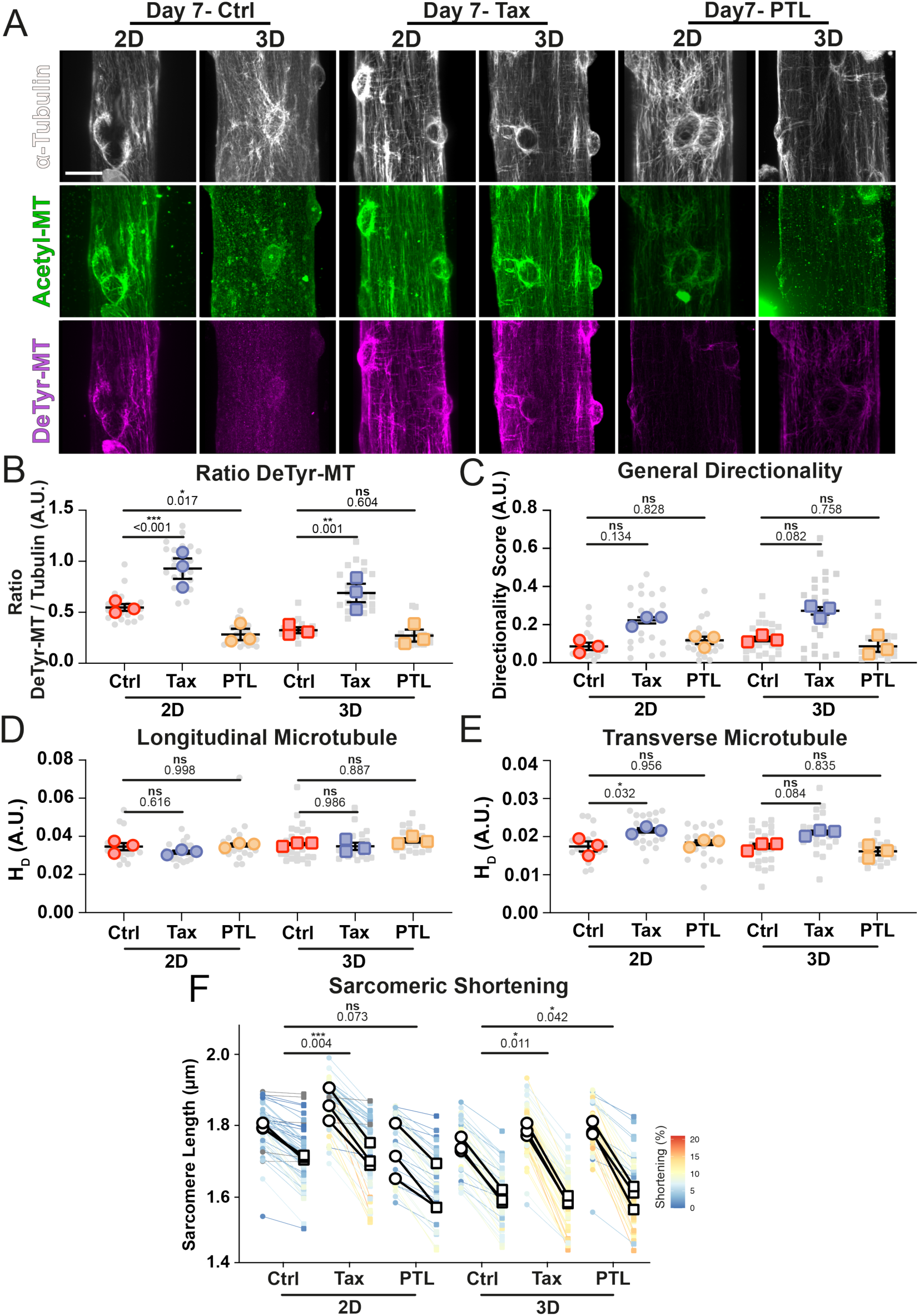
Increasing microtubule organization or inhibition of detyrosinated MT improves muscle fiber contractility. **(A)**: Representative maximum intensity projections of muscle fiber treated with either Taxol (Tax) or parthenolide (PTL) and immunolabeled for α-tubulin (grey), acetylated tubulin (green; acetyl-MT) and detyrosinated tubulin (cyan; deTyr-MT). Scale bar = 20μm. **(B)** Quantification of deTyr-MT fraction in muscle fibers cultured in 2D and 3D on Day 7 and treated with PTL and Tax. **(C)**: Quantification of directionality score as a measure of microtubule lattice network prevalence in muscle fibers cultured in 2D and 3D on Day 7 and treated with PTL and Taxol. **(D)** From the angular histogram extrapolated data to quantify the longitudinal microtubule organization within muscle fibers treated with PTL or Taxol **(E)** From the angular histogram extrapolated data to quantify the transverse microtubule organization within muscle fibers treated with PTL or Taxol **(F)**: The percentage of sarcomere shortening of muscle fibers cultured for 7 days *ex vivo* and treated with PTL and Tax. Data are means ± SEM; *N* = 3 mice and n = muscle fibers. Significance was determined using a linear-mixed model with *p* < 0.05 considered significant with * = *p* < 0.05 and *** = *p* < 0.001.

Taxol produced the opposite effect. It increased deTyr-MT by ∼60% in 2D and ∼110% in 3D (**Figure S10B**), with a smaller rise in acetyl-MT of ∼20% in 2D and ∼50% in 3D (**Figure S10C**). This translated into a marked increase in the deTyr-MT fraction in both conditions, respectively ∼70% in 2D and ∼110% in 3D (**Figure 6B**). The acetyl-MT fraction remained largely unchanged in both 2D and 3D (**Figure S10C**). Although not significant, Taxol trended toward an increase in directionality score, suggesting partial reinforcement of the organized lattice (**Figure 6C**). When MT orientation was further resolved into longitudinal and transverse populations, the longitudinal population was markedly constant across all conditions and pharmacological treatments (**Figure 6D**). However, the transverse MT population remained unchanged when treated with PTL, while Taxol increased the transverse component (**Figure 6E**). These data together suggest that Taxol treatment increases the deTyr-MT levels, while also modulating transverse MT to partially regain the MT-lattice structure.

We next examined whether these changes in MT-PTM, particularly detyrosination, altered contractile output. PTL treatment increased sarcomere shortening by ∼35% in 2D and ∼25% in 3D fibers, relative to their respective control conditions (**Figure 6F & Figure S11**). In 2D, PTL significantly reduced resting sarcomere length without altering sarcomere length at peak contraction, widening the contractile excursion from the resting side. In 3D, where deTyr-MT was already near the detection limit by Day 7 (**Figure 5C**), PTL produced no significant further reduction in the detyrosinated fraction (**Figure S10B**). PTL instead significantly increased resting sarcomere length, opposite to its effect in 2D. Taxol also enhanced contractility (∼45% in 2D, ∼30% in 3D; **Figure 6F & Figure S11**). In 2D, Taxol significantly increased sarcomere length at peak contraction with a parallel, non-significant increase in resting length, whereas in 3D it significantly increased resting length without altering peak. Neither Taxol nor PTL influenced relaxation velocity (**Figure S11D**) in either 2D or 3D, and relaxation time was only different between the PTL and Taxol conditions, with relaxation time taking significantly longer with Taxol treatment compared to PTL.

Together, these data indicate that muscle fiber contractility is governed by two separable determinants of the microtubule network, 1) lattice organization and 2) the degree of deTyr-dependent microtubule–sarcomere coupling, rather than by detyrosination alone (**Figure 7**). Contractile decline over the prolonged culture period was associated with the progressive loss of lattice organization (**Figure 3**). Because the amount of deTyr-MT also decreased over this period, the time course cannot isolate its contribution. This was instead revealed by inhibiting detyrosination with PTL, which, without altering MT abundance or organization (**Figure S10A, 6C**), was sufficient to increase sarcomere shortening (**Figure 6F**), consistent with the higher deTyr fraction and poorer contractility of 2D relative to 3D fibers at Day 7 (**Figure 4E, 5C**). Microtubule organization appears to tune contractility in a separate manner: stabilizing microtubules with Taxol partially restored the transverse lattice and improved shortening (**Figure 6E,F**), accompanied by increased detyrosination, though not driven by it, since PTL lowered detyrosination without changing organization. This suggests that in unloaded muscle fibers, PTL and Taxol improve contractility through distinct routes, so that contractile output reflects how lattice organization and deTyr-dependent coupling jointly set the mechanical relationship between the microtubule network and the sarcomere. Together, our data suggest a model in which microtubule organization and detyrosination act as largely independent regulators of contractility, offering two distinct entry points for preserving or restoring muscle fiber function.

**Figure 7.**
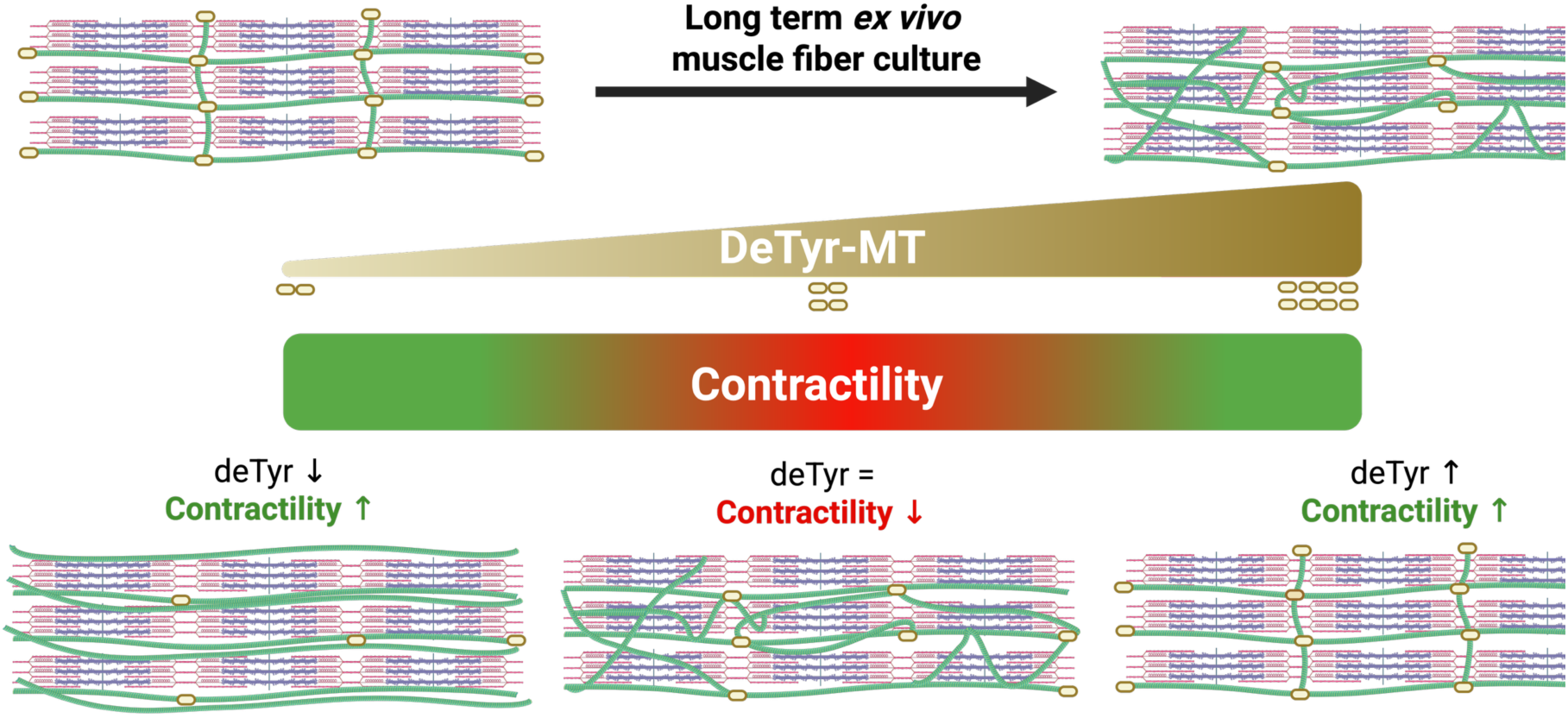
Working model: microtubule organization and detyrosination bidirectionally tune skeletal muscle fiber contractility. Schematic of the proposed relationship between the microtubule (MT) network and sarcomere contractility. *Top:* during long-term *ex vivo* culture under reduced mechanical load, the transverse MT lattice progressively disassembles and the network becomes disorganized and biased toward longitudinally oriented MTs, accompanied by a decline in the detyrosinated-MT (deTyr-MT) fraction. *Middle:* deTyr-MT sets the strength of the indirect MT–sarcomere coupling (springs) that imposes mechanical restraint on the sarcomere; the green-to-red gradient denotes the corresponding contractile state (green, preserved; red, reduced). *Bottom:* contractility is governed by two separable determinants 1) MT lattice organization and 2) deTyr-dependent coupling, so that it can be improved from either direction relative to the disorganized, long-term-cultured state (center; deTyr unchanged, contractility reduced). Lowering the deTyr-MT fraction weakens MT–sarcomere coupling and relieves mechanical restraint, increasing sarcomere shortening without restoring lattice organization. Raising deTyr-MT instead partially restores the transverse lattice, increasing shortening through improved network organization. Contractile output therefore reflects how MT organization and deTyr-dependent coupling jointly set the mechanical relationship between the MT network and the sarcomere.

## Discussion

In this study, we investigated how cytoskeleton remodeling, particularly the MT network, and sarcomere function in skeletal muscle fibers are connected with each other during periods of reduced mechanical activity. Our data identified MT organization and the fraction of deTyr-MTs as key determinants of contractile function in isolated skeletal muscle fibers during prolonged *ex vivo* culture. Isolated muscle fibers undergo a progressive loss of the MT lattice, driven by selective loss of transverse elements that eventually leads to a collapse of the MT lattice. This structural transition in MT-lattice deterioration parallels the decline in sarcomere shortening, supporting a growing body of work showing that skeletal muscle MTs form dynamic and functionally relevant networks that contribute to muscle fiber homeostasis and contractile performance (4,11,29). The present study extends that literature by showing that, in mature muscle fibers, MT geometry and the fraction of deTyr-MT is a stronger predictor of function than MT abundance.

In mature muscle fibers, the MTs form a noncanonical grid-like structure composed of longitudinal and transverse components (19,22,37). This organization has been linked to myofibril patterning and sarcomere assembly during myotube differentiation (38,39). Our findings suggest that this developmental role leaves a durable imprint on adult fiber function, that is, the MT-lattice is not merely a structural hallmark of mature muscle, but a feature required to sustain efficient contraction. The preferential loss of transverse MTs during long-term 2D culture suggests that this network component is sensitive to loss of mechanical and extracellular support. This interpretation is reinforced by the predominance of curved MTs within the transverse networks, consistent with the load-bearing, buckling behavior described for deTyr-MTs in striated muscle (11,27). We also note that curved MT structures frequently coincide with deTyr-MT enriched regions based on qualitative observations, consistent with prior reports linking deTyr-MT arrays to load-bearing and buckling behavior (11,27,28). However, given that contractility can be preserved despite further loss of transverse MTs in 3D culture, these elements alone are unlikely to be the only load-bearing pillars in skeletal muscle fibers. Instead, their contribution may depend on the broader architectural context of the MT-network, although, this relationship was not quantified in this study.

The correlative single-fiber analysis provides strong evidence that MT architecture, not MT abundance, governs contractile performance. At Day 3, when muscle fibers remained heterogenous in both their sarcomere contractility and MT network organization, contractility correlated with MT directionality but not with total MT abundance. This relationship was non-linear and highly informative. Contractility was maintained while directionality stayed high but declined sharply once organization dropped below a threshold value, a behavior consistent with the MT lattice acting as a mechanical network with a minimum organizational requirement. That concept is consistent with the literature on MT-mediated mechanotransduction, where the function of the network depends not simply on polymer load but on how the polymers are arranged and coupled to the contractile apparatus (11,24,40).

Mechanistically, our data align with existing models in which MTs influence contraction through both structural organization and mechanical resistance (11,27,41,42). DeTyr-MTs have been shown to increase cytoskeletal stiffness, induce MT network remodeling, amplify mechanotransduction, and slow contraction by forming a more load-bearing MT network (11,24,27,28,41). In this framework, stabilized MTs are not inherently beneficial, as increased coupling to the sarcomere via desmin can introduce internal resistance to shortening (28). Our pharmacological data support this idea but also suggest an additional layer of regulation involving MT network organization. Reducing detyrosination improved muscle fiber shortening accompanied by a preferential MT-network orientation along the longitudinal axes in 3D culture. This improvement likely reflects reduced internal restraint caused by the elimination of crisscrossing, disorganized MTs. A disorganized MT network can potentially induce physical hindrance, whereas longitudinally aligned MTs may create a smoother path for sarcomere movement (24). This “de-tangling” of the network may allow force to be transmitted more efficiently in the direction of contraction rather than being dispersed by misaligned MT filaments. Conversely, Taxol also improved shortening and partially rescued the MT-lattice. This suggests that reorganizing the MT network rather than strengthening the MT filaments can in some contexts restore cytoskeletal architecture. On one hand, MTs are required for sarcomere assembly and for organizing myosin-containing precursors during differentiation (38,39). On the other hand, stabilized MTs in mature muscle can increase stiffness and mechanical resistance, contributing to impaired force production under pathological conditions (20,23,43).

The 3D culture system provides further insight into this balance. Embedding fibers in laminin-containing matrices preserved viability and maintained contractility far better than 2D culture, yet did not prevent eventual MT lattice breakdown. This suggests that extracellular matrix support can buffer functional decline without fully preserving native MT architecture. One of the functions of the MT-network within muscle fibers is to function as a load bearing scaffold to maintain muscle fiber morphology (44,45). The introduction of laminin-containing matrices likely allowed a physical connection between the extracellular and intracellular environment that shared the experienced mechanical load by the muscle fiber (14). Due to such sharing in the loadbearing capability, the mechanical load is being distributed to both the internal and external compartments, thereby, lowering the internal load bearing demand and allowing remodeling of the cytoskeleton interior (14). Likely, this is being reflected by the weaker MT-sarcomere coupling, rearrangement of the MT filaments, and lower deTyr-MT levels. The literature supports this interpretation, as MT organization in muscle is sensitive to the mechanical milieu and to interactions with sarcolemma and cytoskeletal anchoring systems (14,20,23). Notably, 3D culture biased MTs toward longitudinal alignment rather than maintaining a canonical transverse lattice, indicating that the matrix environment can reshape the network without restoring the native adult pattern. Thus, preserved contractility in 3D does not imply preserved MT architecture; rather, it highlights that muscle function can be sustained by an altered cytoskeletal state as long as the network remains above the threshold required for effective sarcomere performance.

Our data also refine how MT-PTMs should be interpreted in skeletal muscle. Although detyrosination is often viewed as a marker for stable MTs (27,43), its functional impact appears context-dependent. In our system, prolonged culture altered the MT-PTM landscape in parallel with network remodeling, and detyrosination emerged as the more dynamic marker of the functional state of the lattice. The fact that decreasing detyrosination improved contraction indicates that deTyr-MTs contribute to internal mechanical load in this setting, but the simultaneous improvement observed with Taxol suggests that structural stabilization can also be beneficial when it helps maintain a more ordered lattice. Therefore, detyrosination should not be interpreted as uniformly good or bad; its impact depends on whether it reinforces a productive lattice or an over constrained one. One intriguing finding is the effect of MT modulation on resting sarcomere length, as this was the primary mechanism by which sarcomere shortening was enhanced. In freshly isolated cells, acute (2h) treatment with PTL or Taxol does not affect resting sarcomere length (11,30). This discrepancy with our data is likely explained by the fact that our Taxol and PTL experiments were performed after 7 days in culture, a time point where resting sarcomere length has decreased (30). In the 2D, we observed differing roles of Taxol and PTL in terms of their impact on resting sarcomere length, with the former increasing and the latter decreasing it. This effect can be attributed to a non-contractile mechanical function of MTs, which act as compressive structures (27,46,47) and oppose sarcomere shortening at rest. Therefore, we hypothesize that lattice stabilization increases the resting length due to a resistance of the structure to its decrease, while a reduction of deTyr-MT population enables the shortening to the equilibrium set by titin (27,48). Conversely, in 3D the effect of Taxol and PTL was the same, with both resulting in an increase in the resting length. We suspect that in 3D, both treatments have to synergize with an additional resistance to the sarcomere shortening, which comes from the fiber attachment to the matrix, generating passive tension during muscle fiber contraction (49,50). Thus, despite promoting sarcomere shortening in 2D, PTL does not lead to its reduction in 3D due to the mechanical constraint coming from external connections to the matrix. Nevertheless, why both treatments (PTL and Taxol) lead to an increase in sarcomere length in 3D requires further investigation.

The progressive unloading of the muscle fibers of our culture system somewhat mirrors the unloading observed in disuse, and thus, may reflect processes that also occur during muscle unloading *in vivo*. Previous studies have demonstrated that mechanical unloading of skeletal muscle leads to a reduction of α-tubulin protein levels in the predominantly slow-twitch soleus muscle in a load-dependent and reversible manner (32,33). Notably, this response is fiber-type dependent, with α-tubulin content decreasing in slow muscle but not in fast muscle following unloading (33), a distinction that was attributed to the limitations of whole-muscle measurements. Our current results come from isolated muscle fibers from the FDB muscle, which predominantly contains fast-twitch muscle fibers (51). However, it is important to note that previous studies focused on the abundance of tubulin as a means of characterizing the unloading response (32,33). In contrast, our study identifies changes in organization and deTyr-MT level as the key factors that relate to muscle function. We find that reduced load is accompanied by a lower deTyr-MT fraction, consistent with weakened MT–sarcomere coupling (28), and preserved contractility. Conversely, detyrosination increases under conditions of elevated mechanical load and in disease states such as heart failure (27,43), and decreases under the reduced load of our system, suggesting it may act as a bidirectional readout of a muscle cell’s mechanical state. Whether this same detyrosination response contributes to other contexts of reduced muscle activity, such as aging, remains to be examined.

It should be noted that our characterization of the MT network relies primarily on quantitative fluorescence imaging, which carries two detection-related caveats. First, immunofluorescence labeling of fixed muscle fibers reports the polymerized network rather than total tubulin, so our abundance measures reflect only the polymerized MT population rather than total tubulin content (32,33). Nevertheless, our detyrosination data, expressed as a ratio to α-tubulin, are largely insensitive to this. Second, at the very low detyrosination levels reached in 3D fibers by Day 7, we were unable to resolve further reductions, so the absence of a significant PTL effect there likely reflects a detection limit rather than no effect of the treatment. Our assessment of MT–sarcomere coupling is also indirect. The buckling of deTyr-MT under load can be imaged directly in cardiomyocytes (27), but the faster contraction and larger displacements of skeletal muscle fibers make it technically challenging, requiring us to infer coupling from the detyrosination state and organization of the network. Advances in high-speed imaging may eventually allow this to be tested directly.

Finally, our system models reduced mechanical load rather than disuse atrophy *in vivo*, as we see minimal atrophy in this model (30), and our evidence that detyrosination governs contractility is correlative and based on pharmacological interventions rather than genetic, and small-molecule manipulations can act beyond detyrosination alone. Inducible genetic control of detyrosination, for example via a tubulin carboxypeptidase such as vasohibin (52,53), would be required to establish its causal role and clarify how these findings extend to unloading and aging.

Conceptually, our findings support a model in which skeletal muscle contractility is governed by a balance between MT-supported organization and MT-imposed resistance. During healthy maintenance, the transverse MT lattice likely helps preserve sarcomere alignment, intracellular transport, and structural coherence. During prolonged *ex vivo* culture, however, progressive loss of this lattice and changes in PTM state shift the network into a mechanically unfavorable configuration, reducing shortening and slowing relaxation. This provides a mechanistic explanation for why MT organization predicts performance better than MT abundance: what matters is not how much MT remains, but whether the network still retains the spatial order needed to support sarcomere mechanics. The adult fiber MT system is beneficial when it preserves a lattice architecture that supports sarcomere organization and intracellular transport, but becomes detrimental when that same system drifts toward a rigid, maladaptive state that impairs shortening. In this sense, MTs act less like a static scaffold and more like a tunable mechanical regulator.

In summary, our data position MT organization as a functional biomarker and mechanistic regulator of skeletal muscle performance. The adult fiber MT lattice, especially its transverse component, appears to be a critical determinant of contractile competence; its progressive loss during ex vivo culture predicts functional decline more accurately than MT abundance or any single PTM readout. At the same time, the beneficial effects of both reduced detyrosination and Taxol treatment indicate that contractility depends on the balance between lattice order and microtubule-mediated mechanical restraint. This framework reconciles the developmental, homeostatic, and pathological literature on skeletal muscle MTs and suggests that targeting MT architecture may be a powerful strategy to preserve or restore muscle function.

## Acknowledgements

The authors would like to thank Sanna Luijcx, Buram Ardic, Martha Hanzer and Tom Kerkhof for their help with muscle fiber isolation and culture. We would like to thank Sylvia Bogaards for the scientific discussion on ex vivo culture conditions and Dr. Evelyn Ralston for sharing the TeDT code. We would like to thank Michiel Helmes, Valentijn Jansen and Jort van der Geest for their help and expertise in optimizing the Multicell setup and analysis. Lastly, we would like to thank Dr. Diederik Kuster for his helpful discussions during the preparation of the manuscript.

## Funding sources

This study was supported by internal funding from the Department of Physiology, Amsterdam University Medical Center

## Author contributions

O.E. and T.J.K. conceived and designed the study. O.E., E.L., L.V. and N.t.C. performed the experiments and data collection and carried out the formal analysis. O.E. curated the data and produced the figures. T.J.K. provided resources, supervised the project, acquired funding and handled project administration. O.E. and T.J.K. wrote the original draft. E.L., L.V., N.t.C. and T.J.K. reviewed and edited the manuscript. All authors read and approved the final version.

## Competing interests

The authors declare no competing interests.

## Notes

### Competing Interest Statement

The authors have declared no competing interest.

