## Supplemental Figures 1-11 for "Microtubule architecture and detyrosination bidirectionally modulate sarcomere shortening in skeletal muscle fibers"

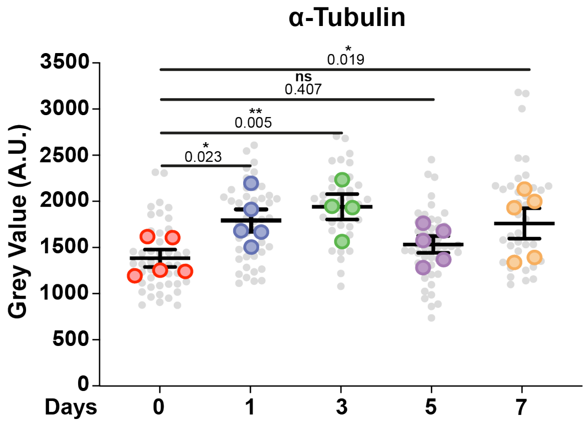


**Figure S1: Effect of culture time on muscle fiber microtubule abundance.** Quantification of the average grey value within each muscle fiber. Data are means ± SEM; $N$ = 5 mice with $n$ = fiber. Significance was determined using a linear-mixed model with $p < 0.05$ considered significant with * = $p < 0.05$ and ** = $p < 0.01$.

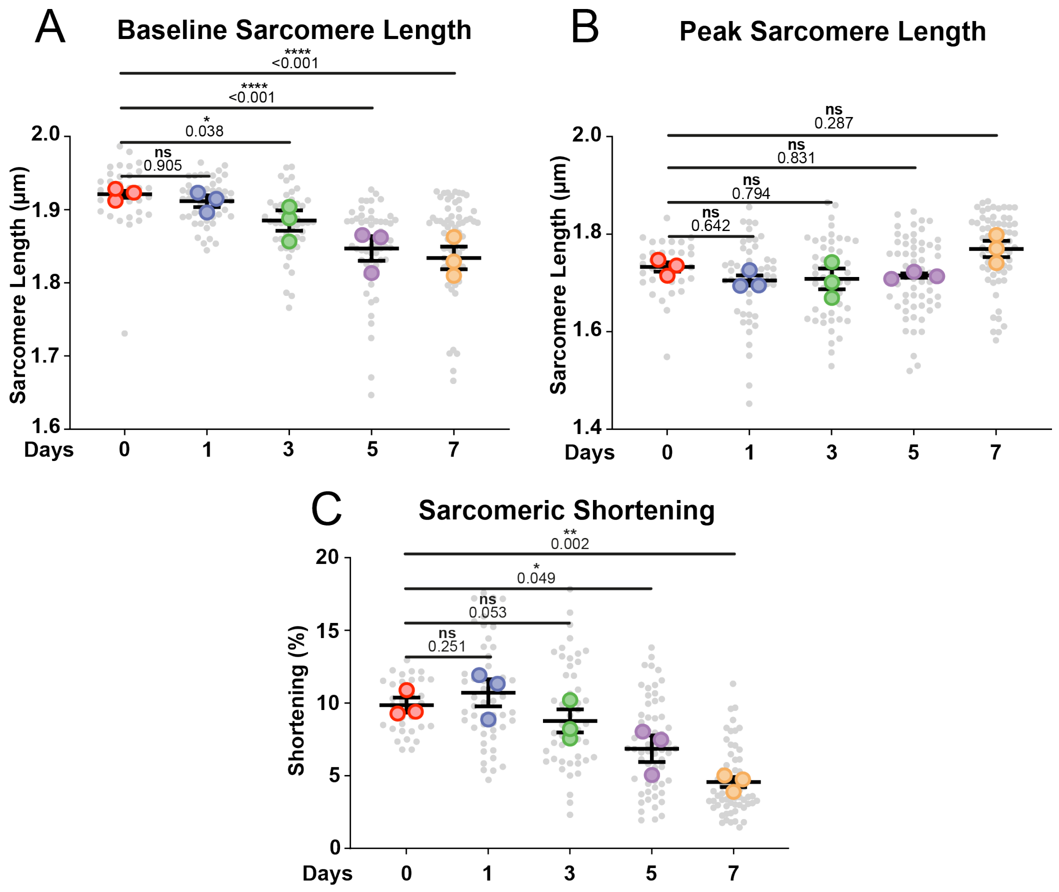


**Figure S2: Contractile measurements of muscle fibers kept in culture on laminin-coated dishes up to 7 days *ex vivo*. (A)** Muscle fiber resting sarcomere length of cultured muscle fibers up to 7 days *ex vivo*. **(B)** Sarcomere lengths during maximal contraction of muscle fibers cultured up to 7 days *ex vivo*. **(C)** The percentage of sarcomere shortening at increasing culture time duration. Data are means ± SEM; $N$ = 3 mice and $n$ = fiber. Significance was determined using a linear-mixed model with $p < 0.05$ considered significant with * = $p < 0.05$, ** = $p < 0.01$, *** = $p < 0.001$, and **** = $p < 0.0001$.

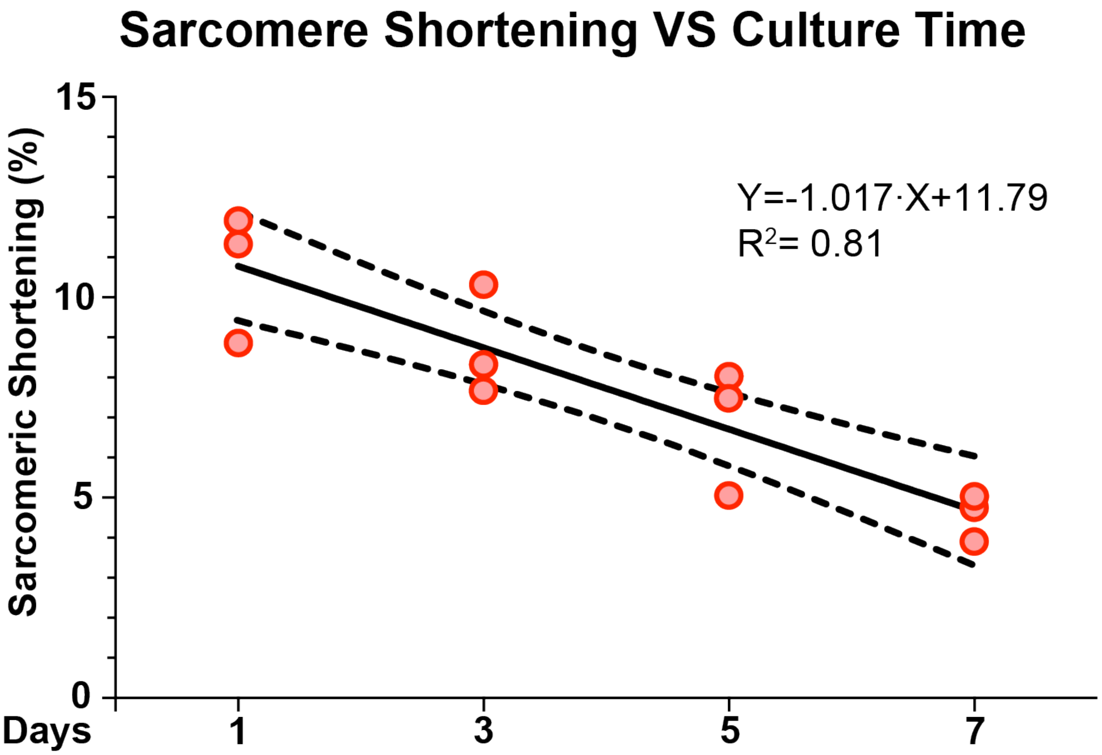


**Figure S3: Correlation between sarcomere shortening and *ex vivo* culture time.** A simple linear regression was fitted through the data points of $N$ = 3 Mice for each time point with a 95% confidence interval.

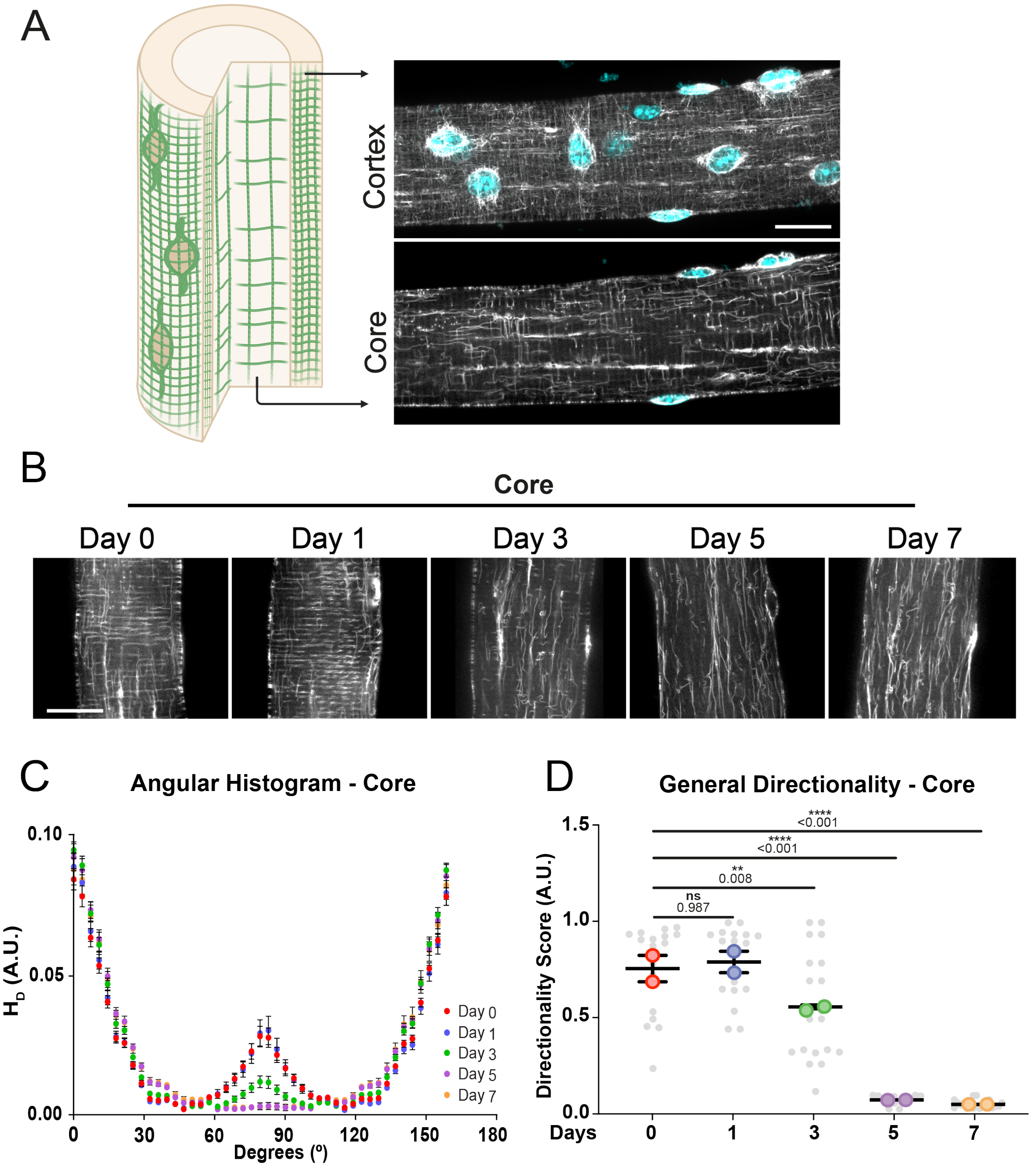


**Figure S4: Validation of the Taxol-based SPY-tubulins neutrality for microtubule remodeling. (A)** Schematic illustration of the dual domain concept composed of a core and cortex region for the microtubules, visualized using SPY-Tubulin. Top right represents maximum intensity projection image while the lower right represents a mid-plane confocal image. All images are stained for microtubule (grey) and nuclei (cyan). Scale bar = 20μm. **(B)** Representative mid-plane confocal images to illustrate the microtubule network in the core of muscle fiber at various time points during *ex vivo* culture immunofluorescently labeled for microtubule (grey). Scale bar = 20μm. **(C)** Angular histogram of the microtubule direction within the core of muscle fibers stained with SPY-Tubulin and cultured up to 7 days *ex vivo*. **(D)**: General directionality score of microtubule organization within the core of muscle fibers cultured up to 7 days *ex vivo*. Data are means ± SEM; $N = 2$ with $n$ = fiber. Significance was determined using a linear-mixed model with $p < 0.05$ considered significant with ** = $p < 0.01$ and **** = $p < 0.0001$.

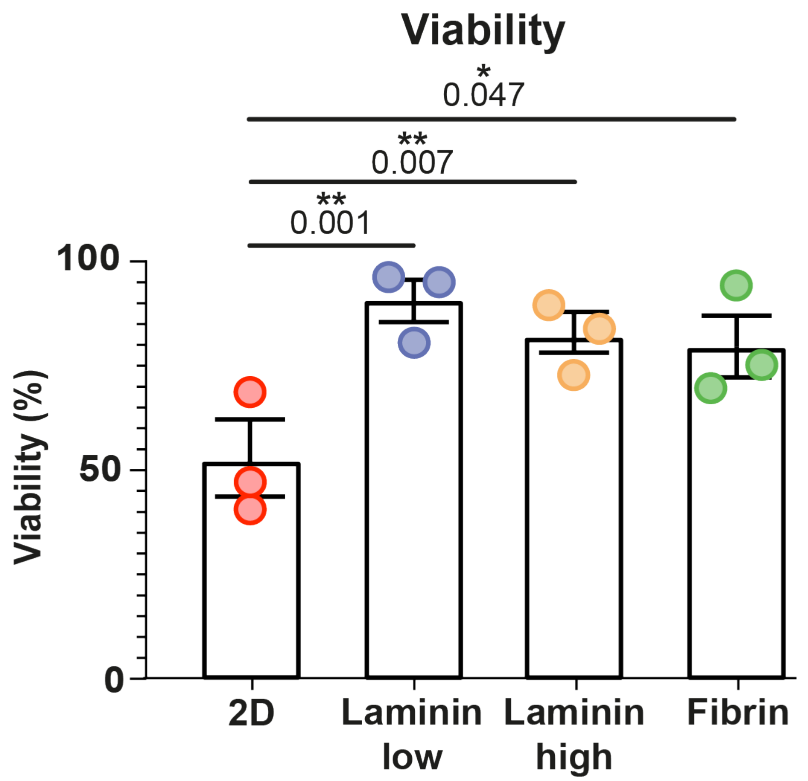


**Figure S5: 3D culture improves muscle fiber viability in *ex vivo* cultured muscle fibers.** Data are means ± SEM; *N* = 3 mice. Significance was determined using a one-way ANOVA with $p < 0.05$ considered significant with * = $p < 0.05$ and ** = $p < 0.01$.

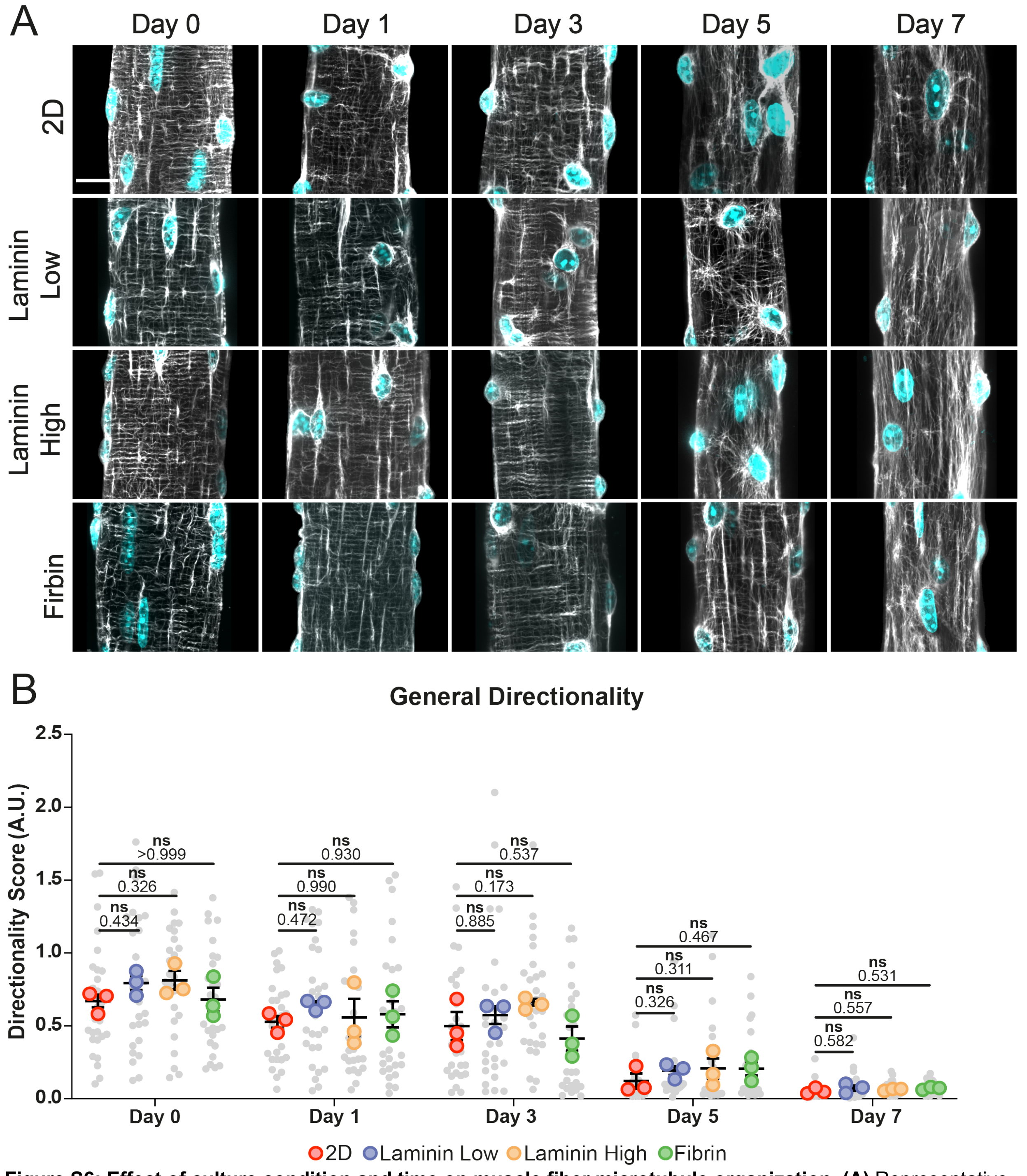


**Figure S6: Effect of culture condition and time on muscle fiber microtubule organization. (A)** Representative confocal maximum intensity projection images taken with a 60X objectives stained for α-tubulin (grey) and nuclei (cyan). Scale bar = 20 μm **(B)** Angular histogram of the preferred microtubule direction within muscle fibers cultured up to 7 days *ex vivo*. Data are means ± SEM; *N* = 3 mice and *n* = fiber. Significance was determined using a linear-mixed model with $p < 0.05$ considered significant.

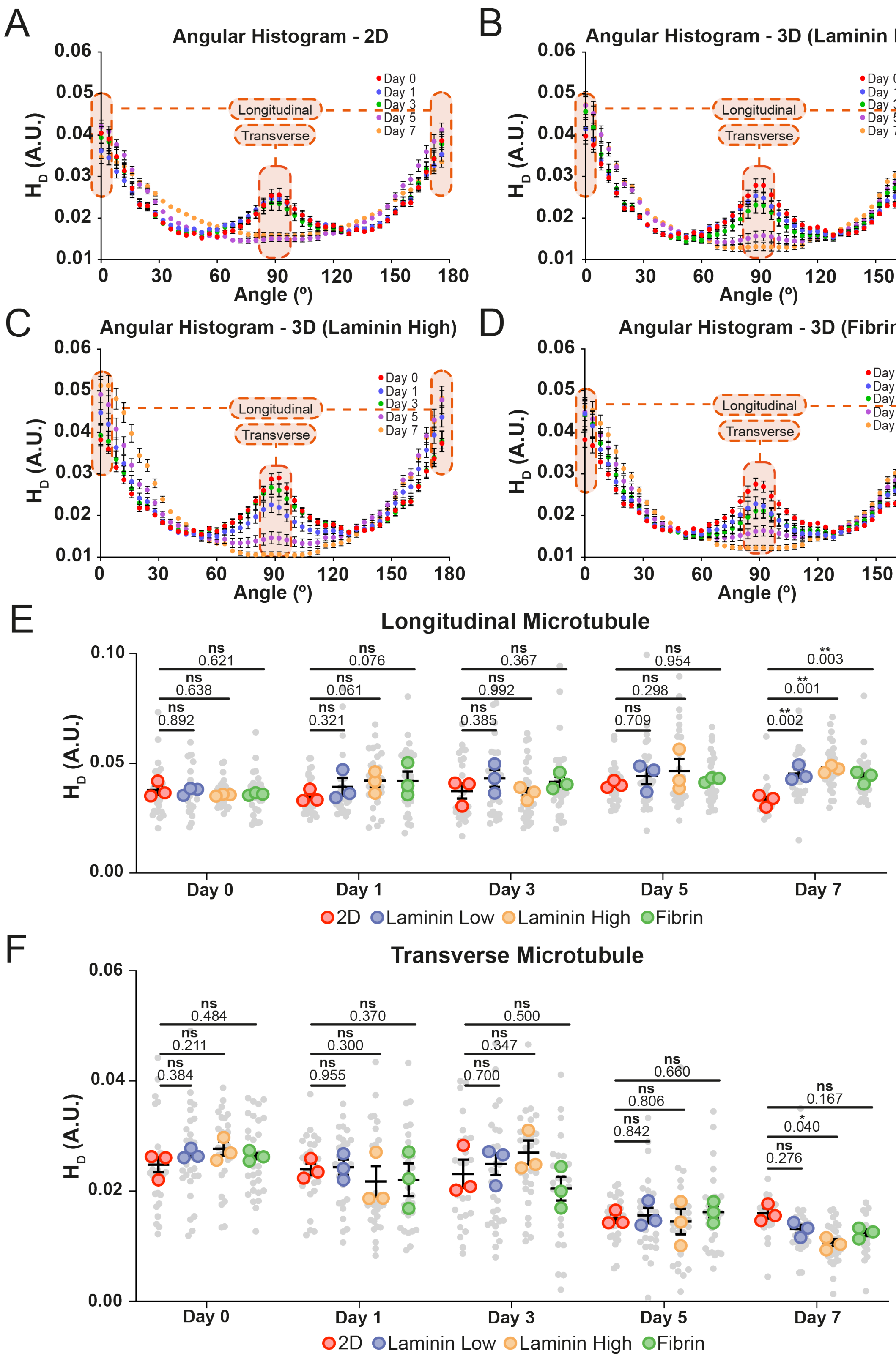


**Figure S7: Effect of culture time on muscle fiber longitudinal and transverse microtubules. (A-D)** Angular histogram of the microtubule direction within muscle fibers cultured up to 7 days *ex vivo* in 2D **(A)**, laminin low gels **(B)**, Laminin high gels **(C)** or fibrin gels **(D)**. **(E)** From the angular histogram extrapolated data to quantify the longitudinal microtubule organization within muscle fibers cultured up to 7 days *ex vivo*. **(F)** From the angular histogram extrapolated data to quantify the transverse microtubule organization within muscle fibers cultured up to 7 days *ex vivo*. Data are means ± SEM; $N$ = 3 mice and $n$ = fiber. Significance was determined using a linear-mixed model with $p < 0.05$ considered significant with * = $p < 0.05$ and ** = $p < 0.01$.

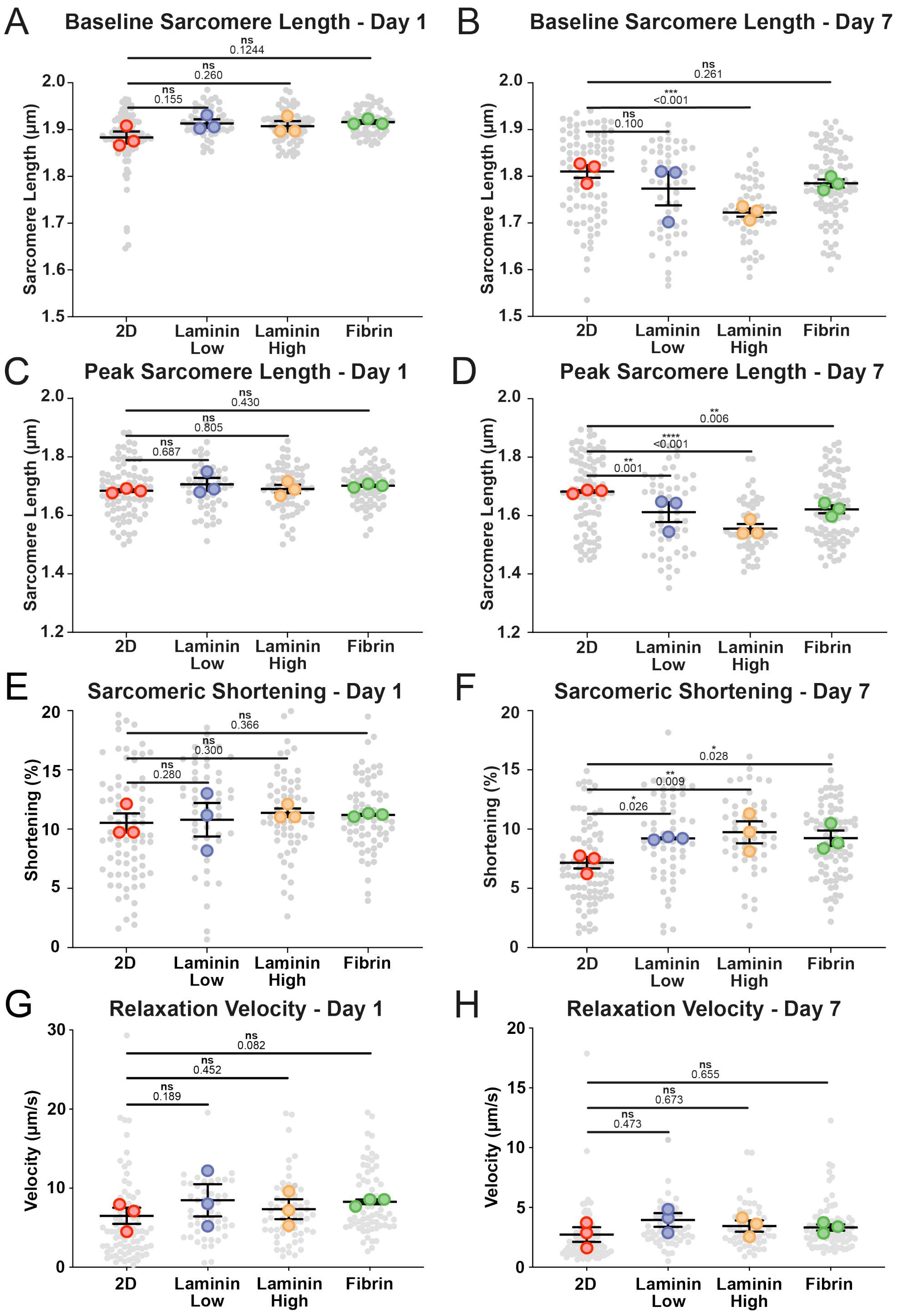


**Figure S8: Embedding fibers in 3D matrices prevents the time-dependent decline in sarcomeric shortening, independent of the laminin concentration. (A,B)** Muscle fiber resting sarcomere length of cultured muscle fibers *ex vivo* on Day 1 **(A)** or Day 7 **(B)**. **(C,D)** Sarcomere length during maximal contraction of muscle fibers cultured *ex vivo* on Day 1 **(C)** or Day 7 **(D)**. **(E,F)** The percentage of sarcomere shortening of muscle fibers cultured *ex vivo* on Day 1 **(E)** or Day 7 **(F)**. **(G,H)** The muscle kinetics measuring their relaxation velocity of muscle fibers cultured *ex vivo* on Day 1 **(G)** or Day 7 **(H)**. Data are means ± SEM; $N$ = 3 mice and $n$ = fiber. Significance was determined using a linear-mixed model with $p < 0.05$ considered significant with * = $p < 0.05$, ** = $p < 0.01$, *** = $p < 0.001$, and **** = $p < 0.0001$.

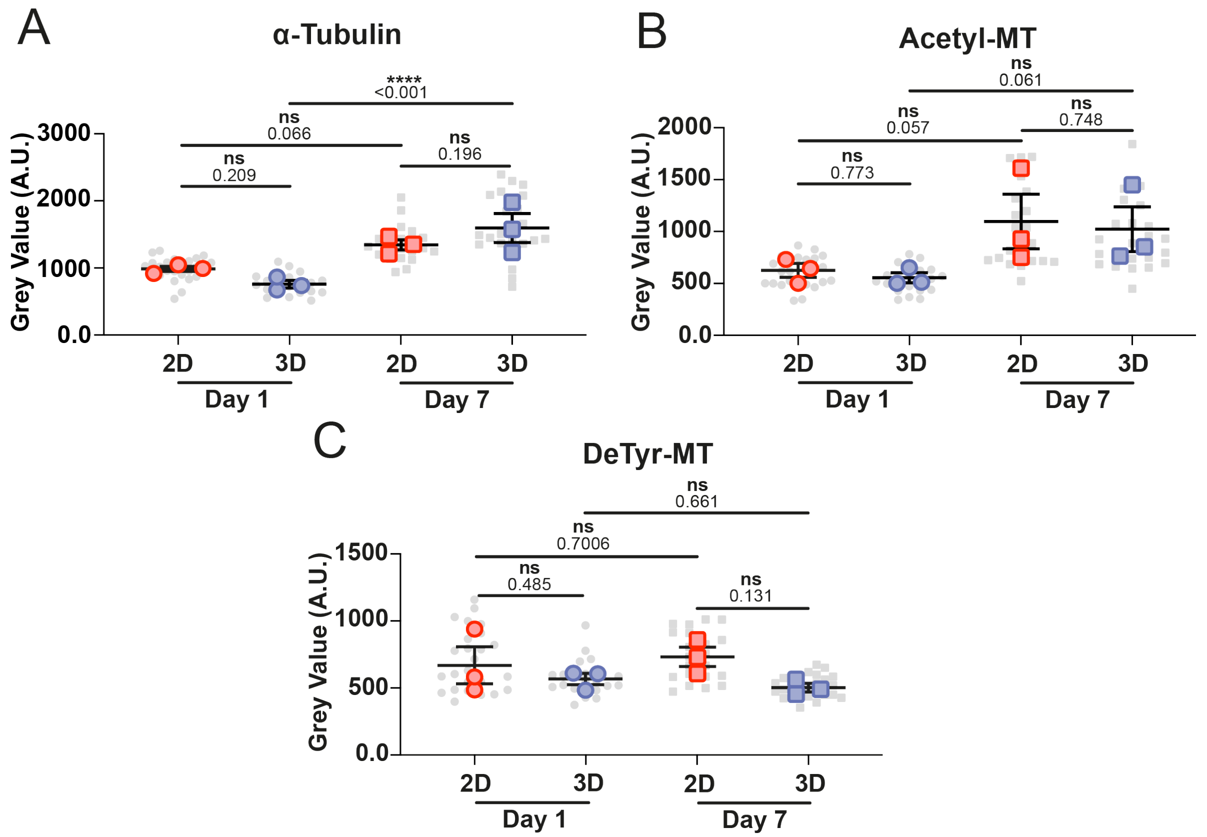


**Figure S9: Effect of culture condition and time on muscle fiber microtubule and posttranslational modification abundance.** (**A**) Quantification of the average grey value of α-tubulin within each muscle fiber cultured in 2D and 3D on Days 1 and 7. (**B**) Quantification of the average grey value of acetylated tubulin (Acetyl-MT) within each muscle fiber cultured in 2D and 3D on Days 1 and 7. (**C**) Quantification of the average grey value of detyrosinated tubulin (DeTyr-MT) within each muscle fiber cultured in 2D and 3D on Days 1 and 7. Data are means ± SEM; N = 3 mice with n = fiber. Significance was determined using a linear-mixed model with $p < 0.05$ considered significant with * = $p < 0.05$ and ** = $p < 0.01$.

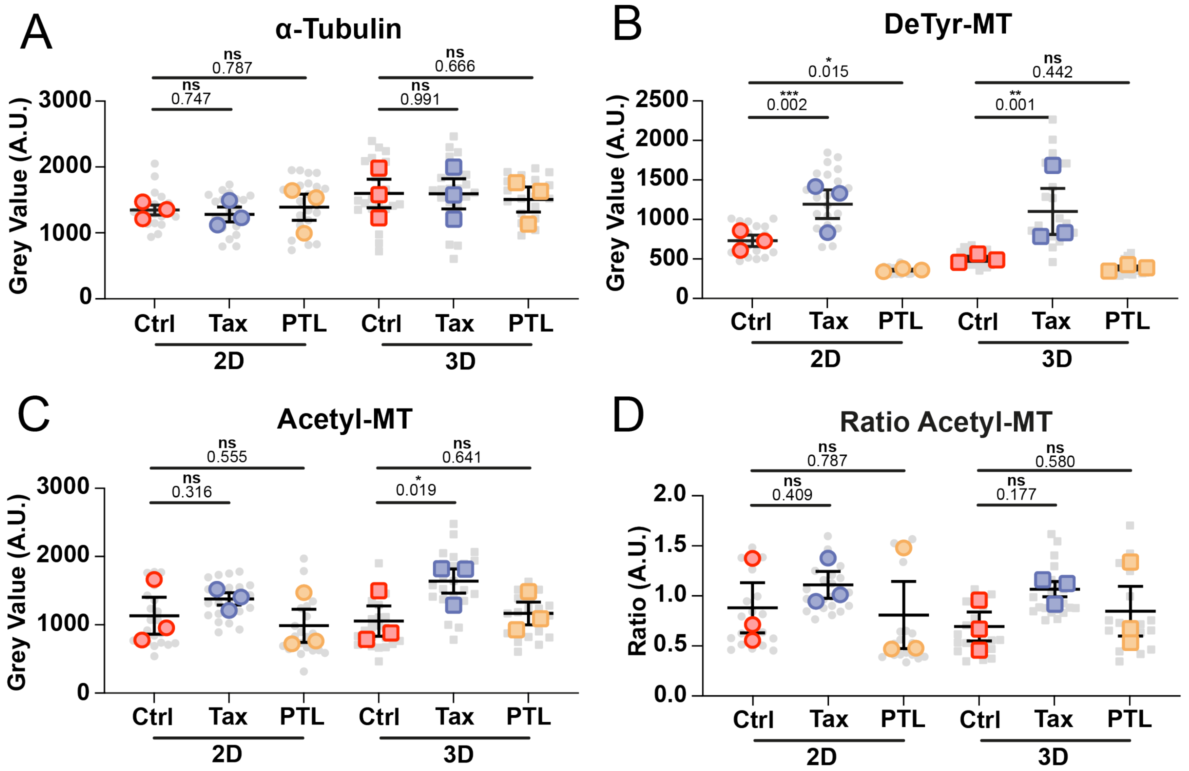


**Figure S10. Quantification of muscle fiber microtubule and posttranslational modification abundance in 2D and 3D in response to Taxol (Tax) or Parthenolide (PTL) treatment.** (**A**) Quantification of the average grey value of α-tubulin within each muscle fiber cultured in 2D and 3D on Day 7 treated with PTL and Tax. (**B**) Quantification of the average grey value of detyrosinated tubulin within each muscle fiber cultured in 2D and 3D on Day 7 treated with PTL and Tax. (**C**) Quantification of the average grey value of acetylated tubulin within each muscle fiber cultured in 2D and 3D on Days 7 treated with PTL and Tax. Data are means ± SEM; N = 3 mice with n = fiber. Significance was determined using a linear-mixed model with $p < 0.05$ considered significant with * = $p < 0.05$ and ** = $p < 0.01$.

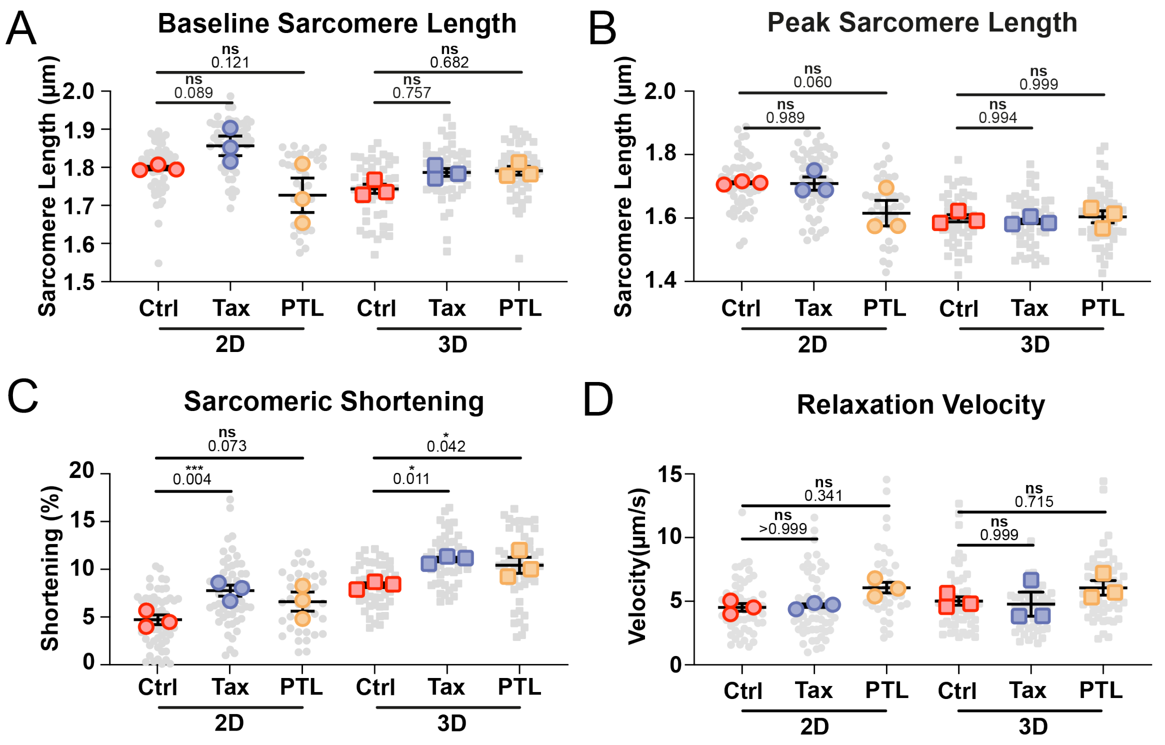


**Figure S11. Quantification of baseline and peak contraction sarcomere length and relaxation kinetics in 2D and 3D in response to Taxol (Tax) or Parthenolide (PTL) treatment**. **(A)** Muscle fiber resting sarcomere length in 2D and 3D on Day 7 treated with Tax or PTL**. (B)** Sarcomere length during maximal contraction of muscle fibers in 2D and 3D on Day 7 treated with Tax or PTL. **(C)** The percentage of sarcomere shortening of muscle fibers in 2D and 3D on Day 7 treated with Tax or PTL. **(D)** The muscle fiber kinetics measuring their relaxation velocity of muscle fibers culture in 2D and 3D on Day 7 treated with Tax or PTL. Data are means ± SEM; N = 3 mice with n = fiber. Significance was determined using a linear-mixed model with $p < 0.05$ considered significant with * = $p < 0.05$ and *** = $p < 0.001$.
